# Computational Insights into How Structural Changes in the Hydrophobic Region of ALS-Linked Mutant SOD1 (G93A) Activate Fluctuations of the Electrostatic Loop: Implications for Aggregation

**DOI:** 10.64898/2026.11.12.766077

**Authors:** Kazuaki Fukushima, Kyoka Yano, Leonard M.G. Chavas, Noriko Fujiwara

## Abstract

Cu/Zn-superoxide dismutase (SOD1) is an essential antioxidant enzyme that converts superoxide radicals into oxygen and hydrogen peroxide. On the other hand, mutations in SOD1 cause familial amyotrophic lateral sclerosis (fALS), and these mutant forms tend to aggregate. Our previous research has shown that residue I35, located in a hydrophobic region, plays a critical role in the intracellular aggregation of SOD1. Furthermore, introducing an additional I35S mutation suppressed aggregate formation and eye degeneration in a Drosophila model caused by ALS-linked mutant SOD1. However, the molecular mechanism linking structural changes induced by ALS mutations to aggregation, and how the compensatory I35S mutation suppresses this aggregation, remain unclear.

To elucidate the relationship between structural changes and aggregation, molecular dynamics simulations were performed on three types of SOD1 monomers: wild type (WT), G93A (an fALS mutant), and the G93A/I35S double mutant. In WT, the metal-binding loop (loop IV) exhibited slightly larger fluctuations, whereas in G93A, the electrostatic loop (loop VII) fluctuates significantly more. Detailed analysis revealed that the G93A mutation shifts the hydrogen-bonding partner of the 93rd residue from L38 in WT to G37 in G93A. This change destabilizes the V119-L144 hydrogen bond, thereby enhancing fluctuations in β-strand 8 and the electrostatic loop. Notably, the I35S mutation in the G93A background restores the WT-like packing of the hydrophobic core and reestablishes the V119-L144 hydrogen bond, reducing the abnormal electrostatic loop fluctuations.

These findings provide a mechanistic link between a single-point mutation and conformational changes involved in SOD1 aggregation. Our results also suggest that targeting the hydrophobic region may represent a novel therapeutic strategy for SOD1-linked fALS.

## Introduction

Cu/Zn-superoxide dismutase (SOD1) is a homodimer enzyme that plays a critical role in cellular defense against oxidative stress. The enzyme contains one zinc and one copper ion *per* subunit; each composed of 153 amino acid residues. Primarily located in the cytosol, SOD1 catalyzes the conversion of the superoxide radical into hydrogen peroxide and molecular oxygen *via* the Cu^+^/Cu^2+^ redox cycle in the active site [1, 2]. Despite its essential role as an antioxidative enzyme, mutations in SOD1 have been implicated in familial amyotrophic lateral sclerosis (fALS) [3–5].

ALS is a progressive neurodegenerative disease affecting motor neurons in the brain and spinal cord, with its underlying causes remaining largely unclear. Transgenic mouse models have shown that overexpression of human ALS-linked mutant SOD1 results in the development of ALS-like symptoms, whereas SOD1 knockout mice do not exhibit such phenotypes [6]. These observations have spurred numerous comparative studies of recombinant wild-type (WT) and ALS mutant SOD1 proteins. Structurally, SOD1 is characterized by a β-barrel structure composed of eight β-strands connected by seven loops [7, 8]. Among these loops, loop IV (the metal-binding loop) and loop VII (the electrostatic loop) are intimately associated with both the catalytic activity of the enzyme and its aggregation process [9–11]. Although WT SOD1 is inherently stable, ALS-linked mutations compromise its stability, rendering the protein more susceptible to denaturation and aggregation [12, 13]. A prevailing hypothesis is that destabilization of the electrostatic loop is a common characteristic of ALS-associated mutants [14]. The destabilization may lead to conformational changes in the metal-binding loop, resulting in the loss of Zn^2+^ ions and subsequent protein denaturation [15, 16]. Studies on recombinant SOD1 aggregation suggested that the process involves (i) dissociation of the native homodimer into monomers, (ii) reduction of the intramolecular disulfide bond (C57–C146), and (iii) loss of metal ions [17–20]. In addition, pH and salt concentration are known to influence aggregation [21].

We previously reported that residue I35 in the hydrophobic region plays a critical role in the intracellular aggregation of SOD1. In that work, analyses using BioID2 and EGFP fusion expression systems revealed that mutating I35 to S35 markedly suppresses the formation of intracellular aggregates of the G93A mutant. (Fig. S1) [22]. Moreover, McAlary *et al*. reported that introducing an I35P mutation in the G85R mutant also suppresses intracellular aggregation [23]. Furthermore, we recently reported that the neurodegeneration observed in the compound eyes of adult flies expressing BioID2-G93A-EGFP is suppressed by the additional I35S mutation (G93A/I35S) [24]. These findings suggest that subtle structural changes in the hydrophobic region including I35 may have long-range effects that promote aggregation and toxicity in ALS-mutant SOD1. We therefore hypothesized the possibility that subtle structural changes in this hydrophobic region were responsible for the destabilization of the electrostatic loop caused by the G93A mutation. Furthermore, we speculated that introducing the I35S mutation might suppress the destabilization in this electrostatic loop. MD simulation is a powerful technique used to study temporal changes in the positions of atoms under defined conditions, and its applications in molecular biology have expanded widely [25–28]. This method allows us to visualize interactions and motions of biopolymers such as proteins, thereby enabling us to assess how overall structures and specific fluctuations respond to partial mutations or ligand binding. Over recent years, several MD studies have been performed on SOD1 to explore its stability and aggregation mechanisms [29–44]. For example, Souza et al. identified a transmission pathway mediated by hydrogen bonds between multiple residues that explains the effect of the G93A mutation on the electrostatic loop and revealed that the opening of this loop leads to the dissociation of Zn^2+^ ions [29]. In addition, Zhu *et al*. applied a neural relational inference model to MD simulations to uncover long-range allosteric interactions within proteins, and they identified an allosteric pathway in the G93A mutant that operates alongside hydrogen-bond interactions [44]. However, the molecular mechanism linking these small steric changes to altered SOD1 conformation and intracellular aggregation remains to be elucidated.

To study the relationship between conformational change and aggregation, we first conducted extensive 300-ns atomistic molecular dynamics (MD) simulations on SOD1 monomers of WT, the fALS-associated G93A mutant. Based on the differences observed, we subsequently performed similar MD simulations on the G93A/I35S double mutant, which is resistant to aggregation. Although SOD1 functions as a dimer *in vivo*, previous studies indicate that dissociation into monomers is a necessary preliminary step for aggregation [3]. Therefore, focusing on the monomer allows us to capture the early conformational changes that underlie aggregation.

This work aims to clarify the allosteric mechanism by which a single-point mutation SOD1 can induce long-range conformational changes that ultimately lead to aggregation, a key event in the pathogenesis of ALS. In doing so, our study provides insights that may help guide the development of targeted therapeutic strategies aimed at stabilizing the hydrophobic core of SOD1.

## Results

### The fluctuations of the SOD1 molecule are large in the metal-binding loop in WT, and large in the electrostatic loop in G93A

Figure 1A shows the geometry of the WT SOD1 dimer, and Figure 1B illustrates the monomer geometry with the names of the loops and β-strands. Notably, the metal-binding loop (loop IV, residues 49-82) and the electrostatic loop (loop VII, residues 121-142) are highlighted in green and blue, respectively. Figure 1C identifies the sites associated with the structural changes in G93A centered on the hydrophobic region.

**Figure 1.**
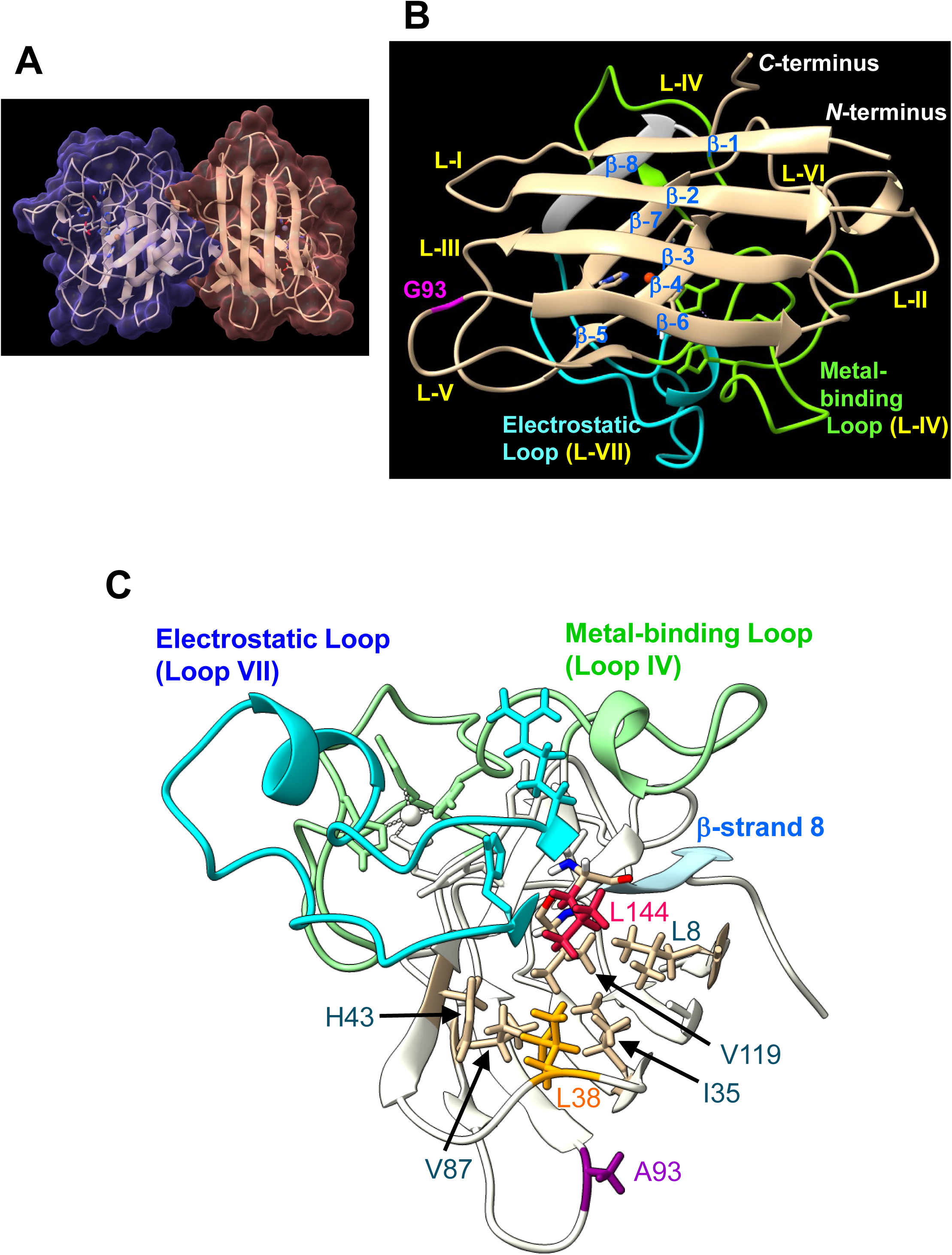
Geometry of the wild type SOD1 and G93A mutant. ***A***, Dimer geometry (PDB ID: 3T5W). ***B***, Monomer geometry. The β-*n* and the L-*m* represent β-sheet and loop, respectively. ***C***, Hydrophobic region of G93A. The steric influence of A93 is transmitted from L38 to β-strand 8 and the electrostatic loop via the hydrophobic region (L8, I35, H43, V87, V119, L144).

We first performed five independent 300-ns MD simulations for both WT and G93A SOD1 monomers. RMSD calculations for WT and G93A yielded different pattern profiles, indicating that distinct structural changes occurred in each production run (Fig. S2). In all runs, the RMSD from the initial structure remained within approximately 0.2 nm (2.0 Å), which confirms that no significant structural changes or denaturation events occurred during the 300-ns simulations, with the overall globular structure of SOD1 that was maintained for both proteins. Thus, we proceeded to analyze which regions of the WT and G93A proteins exhibited increased fluctuations.

Root-mean-square fluctuation (RMSF) analysis for each run revealed that both WT (Figs. S3-1) and G93A (Fig. S3-2) displayed large fluctuations in the metal-binding loop (residues 49–82) and the electrostatic loop (residues 121–142). However, in WT, prominent fluctuations were observed in the metal-binding loop in Runs 1 and 4 (Fig. S3-1). By contrast, G93A exhibited a more heterogeneous pattern: large fluctuations in the electrostatic loop were evident in Runs 2 and 3, while Run 5 showed prominent fluctuations in the metal-binding loop, and Run 4 revealed significant fluctuation in loop V that contains A93 (Fig. S3-2). When averaging the RMSF over all runs, we found that both proteins showed large fluctuations in the metal-binding loop (particularly residues 49–62, with T54 as the largest fluctuating residue) and in the electrostatic loop (residues 128–135, with G130 as the largest fluctuating residue) (Fig. 2A). However, compared with WT, the G93A mutant exhibited notably enhanced fluctuations in the electrostatic loop (Loop VII), while the fluctuations of the metal-binding loop (Loop IV) were reduced (Table S3-1).

**Figure 2.**
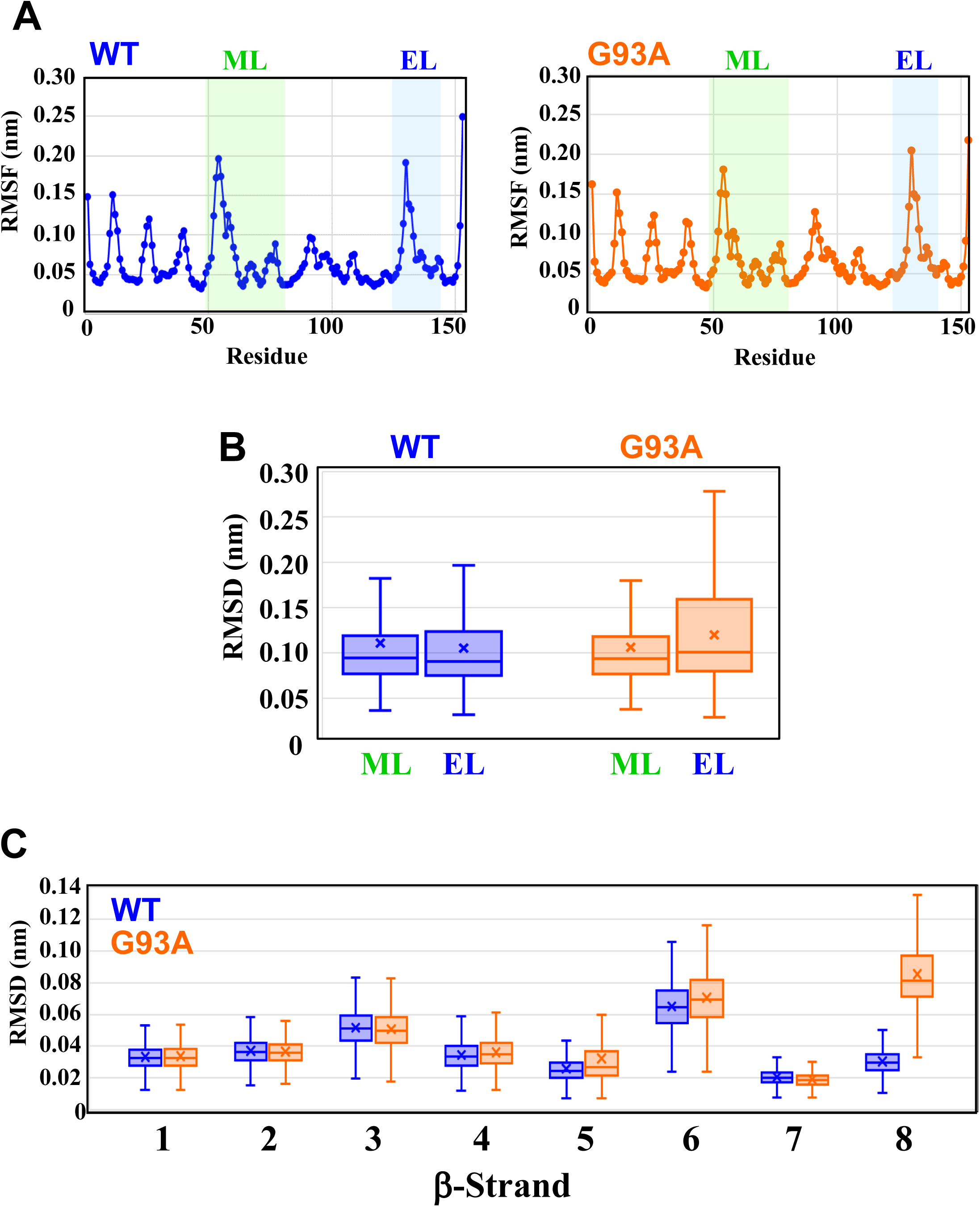
Fluctuation of WT and G93A. ***A***, RMSF of WT and G93A. The metal-binding loop (ML: 49-82) is indicated by the green ribbon, and the electrostatic loop (EL: 121-142) is indicated by the blue ribbon at the bottom. ***B***, Box-and-whisker plot showing the distribution of RMSD of the metal-binding loop (ML) and the electrostatic loop (EL) in all geometries (*n*=150,000) from Run 1 to Run 5. The box represents IQR (interquartile range: 25-75th percentile), the horizontal line inside the box represents the median value, and the cross mark represents the average value. The top of the whiskers above the box represents the maximum value, and that below the bottom represents the minimum value. WT is drawn in blue and G93A is drawn in orange. ***C***, Box-and-whisker plot showing the distribution of RMSD of the β-strands defined by ChimeraX 1.5 in all geometries (*n*=150,000) from Run 1 to Run 5. The β-strands 1 to 8 are represented by 1-8 (1: 2-8, 2: 15-22, 3: 29-36, 4: 41-48, 5: 86-89, 6: 95-101, 7:116-120, 8: 144-149). The box represents IQR (interquartile range: 25-75th percentile), the horizontal line inside the box represents the median value, and the cross mark represents the average value. The top of the whiskers above the box represents the maximum value, and that below the bottom represents the minimum value. WT is drawn in blue and G93A is drawn in orange.

Since the RMSF analysis indicated that WT is predisposed to sampling different conformations in the metal-binding loop while the G93A mutant favors alternate conformations in the electrostatic loop, we performed RMSD calculations on these loops by fitting the backbone atoms (C, C_α_, and N) of both proteins. As shown in Figure 2B, the box-and-whisker plot of the RMSD distribution indicates that the fluctuations of the metal-binding loop are slightly larger than those of the electrostatic loop in WT, whereas in G93A the electrostatic loop exhibits remarkably larger fluctuations compared to the metal-binding loop. In addition, the overall fluctuations in the electrostatic loop are clearly greater in G93A than in WT (Fig. 2B). A closer examination at the RMSD time courses of the electrostatic loop in WT revealed that, while Runs 1–3 and 5 exhibited only small fluctuations, Run 4 displayed relatively large and sustained fluctuations (Fig. S4-1, left). In contrast, G93A demonstrated large fluctuations in Runs 2 and 3 and long-term fluctuations in Run 5 (Fig. S4-1, right). However, RMSD time courses of the metal-binding loop revealed that WT demonstrated large fluctuations in Runs 1 and 4 while G93A exhibited relatively large fluctuations only in Run 5 (Fig. S4-2). Although the box plot for the metal-binding loop in Figure 2B shows no clear difference between WT and G93A, the statistical values presented in Table S4-1 indicate that the fluctuations of the metal-binding loop in WT are, on average, greater than that in G93A.

To further clarify the relationship between fluctuations in the electrostatic and metal-binding loops and the changes occurring in the protein backbone, we compared the RMSD of all eight β-strands in WT and G93A. As shown in Figure 2C, the RMSD of β-strand 8 of G93A was significantly larger than that of the other β-strands whereas the RMSD of β-strand 8 in WT remained low. The time course of RMSD for β-strand 8 (Fig. S5) also indicates that its fluctuations are larger in G93A compared to WT. Since β-strand 8 is directly connected to the electrostatic loop (loop VII), this observation suggests that the enhanced fluctuation of β-strand 8 in G93A is closely related to the increased mobility of the electrostatic loop.

A comparison of the 2D-plot of free energy landscapes obtained from principal component analysis (PCA) revealed that while the WT exhibited a single stable structure, the G93A possessed two stable structures (Fig. 3A). Although the stable structure of WT (WT-1) was similar to one of the stable structures of G93A (G93A-1), the two stable structures of the G93A (G93A-1 and G93A-2) exhibited large different structural conformations in the electrostatic loop region (Fig. 3B). These results suggest that G93A adopts a second stable conformation characterized by an electrostatic loop conformation different from that of WT, and the larger fluctuations in the electrostatic loop and β-strand 8 in G93A (Figs. 2B and 2C) are thought to be attributable to this structural difference.

**Figure 3.**
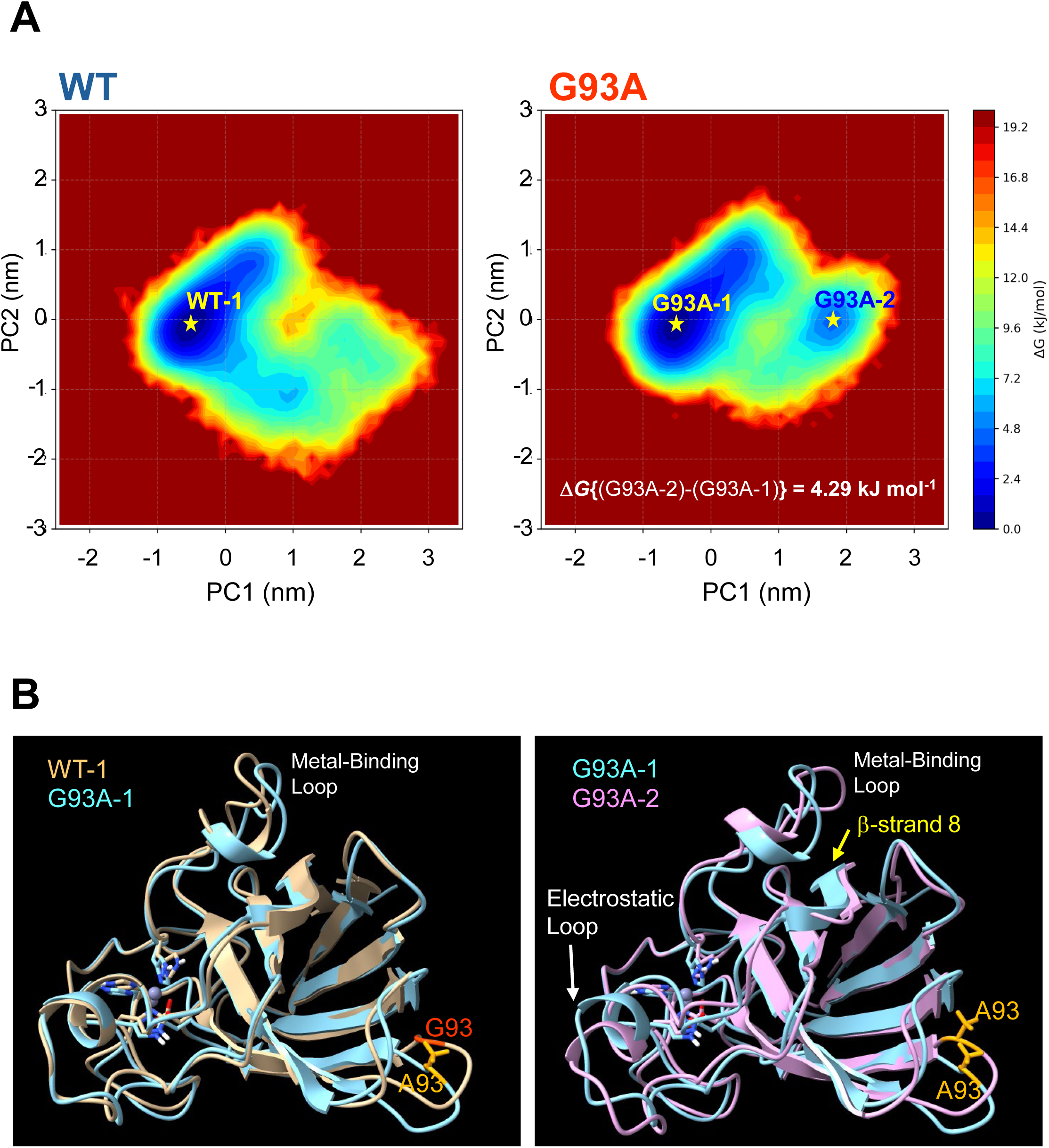
2D-Plot of Free Energy Landscape. ***A***, Free energy landscapes (FELs) of WT (left) and G93A (right) as a function of the first (PC1) and second (PC2) principal components. Principal component analysis (PCA) was performed on the combined backbone trajectory of both systems. Color gradients indicate relative free energy (Δ*G*), where lower-energy basins correspond to conformational states with higher probability densities. ***B***, Structural comparisons of the free energy minima. Left: Superposition of the most stable conformation of WT (WT-1, brown) and that of G93A (G93A-1, cyan). Right: Superposition of the most stable conformation (G93A-1, cyan) and the second most stable conformation (G93A-2, magenta) within the G93A trajectory.

### Mechanism by which the G93A Mutation Causes Changes in the Structural Fluctuation Pattern

#### 1. Mutation from G93 to A93 changes the distances between L38 and some amino acids in the hydrophobic region

We next investigated the mechanism by which the single point mutation from WT to G93A leads to structural changes in the electrostatic loop and β-strand 8. Several studies have reported that the effect of changing G93 to A93 appears as a change in the position of the spatially adjacent L38 [23, 44]. However, the detailed interactions among the amino acids altered by the G93A mutation remain unclear. To investigate this, we first analyzed the interatomic distances between G93 (or A93) and L38 or G37. In WT, the distances between the C=O group of G93 and the NH group of L38 exhibited a sharp distribution with a peak at 0.205 nm over for the five production runs (Fig. 4A(a)). In contrast, in G93A the distances between the C=O group of A93 and the NH group of G37 showed a sharp distribution with a peak at 0.200 nm (Fig. 4A(b)). These results indicate that the hydrogen-bond partner of the C=O group of G93 (or A93) changes from L38 in WT to G37 in G93A. The model depicted in Figure 4B suggests that the steric hindrance introduced by the methyl group of A93 is responsible for this shift in hydrogen-bonding.

**Figure 4.**
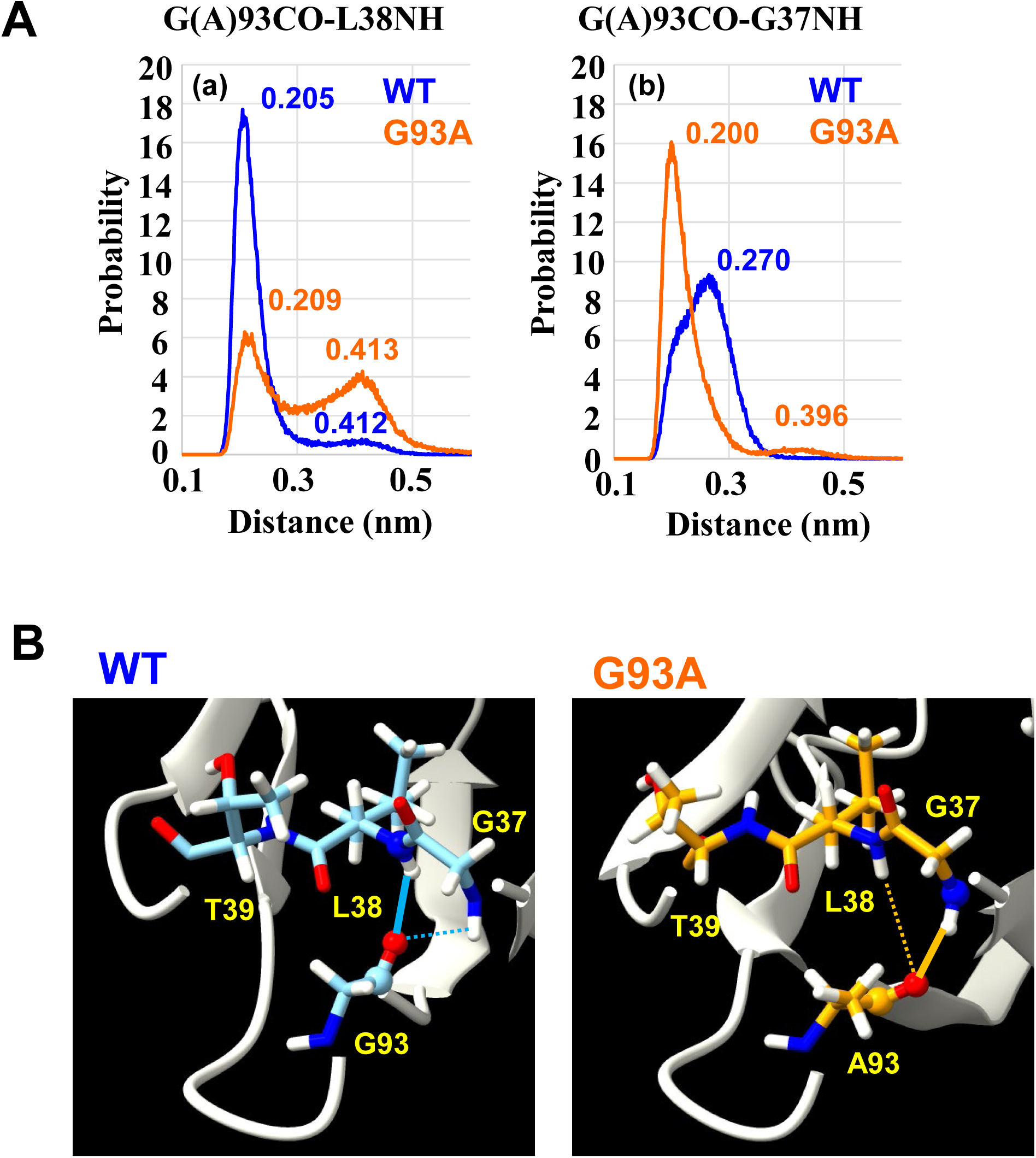
Change in Hydrogen-Bonding Pair between G(A)93 and G37 or L38. ***A***, (a), Distribution of the hydrogen-bond distances between the C=O group of G(A)93 and the N-H group of L38 in all geometries (*n*=150,000) from Run 1 to Run 5. (b), Distribution of the hydrogen-bond distances between the C=O group of G(A)93 and the N-H group of G37 in all geometries (*n*=150,000) from Run 1 to Run 5. WT is drawn in blue and G93A is drawn in orange in (a) and (b). The peak apex is indicated in nm. ***B***, Three-dimensional arrangement of G93 and the residues spatially close to G93 in WT, and that of A93 and the residues spatially close to A93 in G93A (G37, L38, and T39). The geometries are those of the cluster 1 of Run 1 in WT and G93A.

The change in the hydrogen-bond pair, from G93-L38 in WT to A93-G37 in G93A, also affects the distance distribution between the α-carbon (C_α_) of G93 (or A93) and those of spatially adjacent residues, namely G37, L38, and T39 in loop III. As shown in Figure S6-1, the distances between the C_α_ of G93 (or A93) and the C_α_ of G37 were nearly constant in both WT and G93A, whereas the peak for the distances between the C_α_ of A93 and the C_α_ of L38 in G93A (0.528 nm) was approximately 0.08 nm longer than the corresponding distances between L38 and G93 in WT (0.449 nm). This indicates that the distance between G93 (or A93) and L38 increases as a consequence of the altered hydrogen-bond pairing. Moreover, the distances between the C_α_ of G93 (or A93) and the C_α_ of T39 in G93A display two distinct peaks (Fig. S6-1). These two peaks are attributed to the inherent flexibility of loop III, which allows T39 to occupy two positions (Fig. S6-2). Because T39 is capable of forming intermolecular hydrogen bonds with neighboring residues, changes in its position can significantly affect the overall structure of the molecule, a point that will be discussed further below.

L38 is significantly affected in its conformation by the mutation from G93 to A93 (Fig. 4). Since the hydrophobic side chain of L38 is located within the interior of the protein (Fig. 1C), even a slight change in its position can induce conformational alterations in surrounding residues, including I35, H43, V87, V119, and L144. When comparing the distances between the C_α_ of L38 and the C_α_ of these adjacent residues, we found that the distances of L38–I35 (Fig. 5A(a)) and L38–V87 (Fig. 5A(b)) were shorter in G93A than in WT. Conversely, the distance of L38–L144 (Fig. 5A(c)) was longer in G93A compared to WT. In G93A, the mutation of G93 to A93 results in a repositioning of L38 toward the exterior of the protein relative to WT (Fig. 5B). This outward movement of L38 causes L144 to shift upward, while V87 and I35 move inward and assume positions closer to each other. (Note that Figure 4B presents L38 and its surrounding residues from a different angle than that in Figure 1C.)

**Figure 5.**
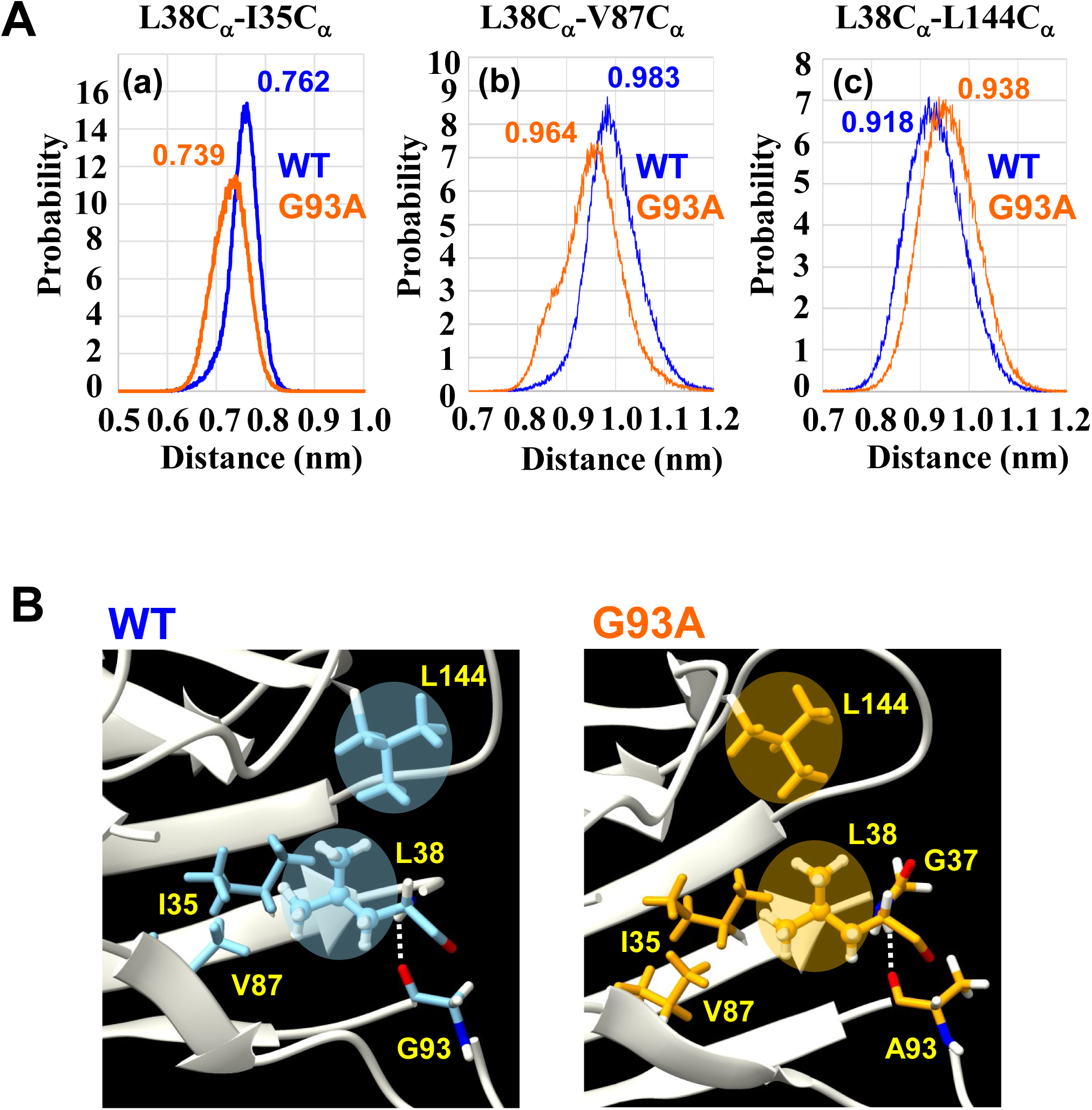
Geometry Change around L38 and β-Strand 8 in WT and G93A. ***A***, Distribution of distances between C_α_ of L38 and C_α_ of the residues spatially close to L38 in all geometries (*n*=150,000) from Run 1 to Run 5. (a), C_α_ of L38 and C_α_ of I35. (b), C_α_ of L38 and C_α_ of L144. (c), C_α_ of L38 and C_α_ of V87. WT is drawn in blue and G93A is drawn in orange. The peak apex is indicated in nm. ***B***, Three-dimensional arrangement of L38 and the residues spatially close to L38 in WT and in G93A (G(A)93, I35, G37, V87, and L144). The right side is outside of the protein, and the left side is inside of the protein. The geometries are those of the cluster 1 of Run 1 in WT and G93A. ***C,*** Three-dimensional arrangement of L8 and β-strand 8 in WT and in G93A. Hγ of L8 is drawn in magenta. The geometries are those of the cluster 1 of Run 1 in WT and G93A.

#### 2. The steric change in L38 reduces the V119-L144 hydrogen-bond

The steric changes in L38 affect the distribution of L38-L144 distances (Fig. 5A(c)), which in turn is expected to influence the V119-L144 hydrogen bond. Figure 6A displays the distribution of the O^…^H distances (a) and O^…^H-N angles (b) between the C=O group of V119 and the NH group of L144 in WT and G93A. In WT, the O^…^H distances are frequently less than 0.2 nm and the O^…^H-N angles are often less than 30°, indicating that the C=O group of V119 and the NH group of L144 are well oriented for hydrogen-bond formation (Fig. 6B, left). By contrast, in G93A the O^…^H distances are generally longer and the O^…^H-N angles are larger, resulting in conformations that are not favorable for hydrogen-bond formation (Fig. 6B, right). These differences in distances and angles are likely caused by the altered position of L144 between G93A and WT, and this change is expected to affect the geometry of the electrostatic loop that is adjacent to L144.

**Figure 6.**
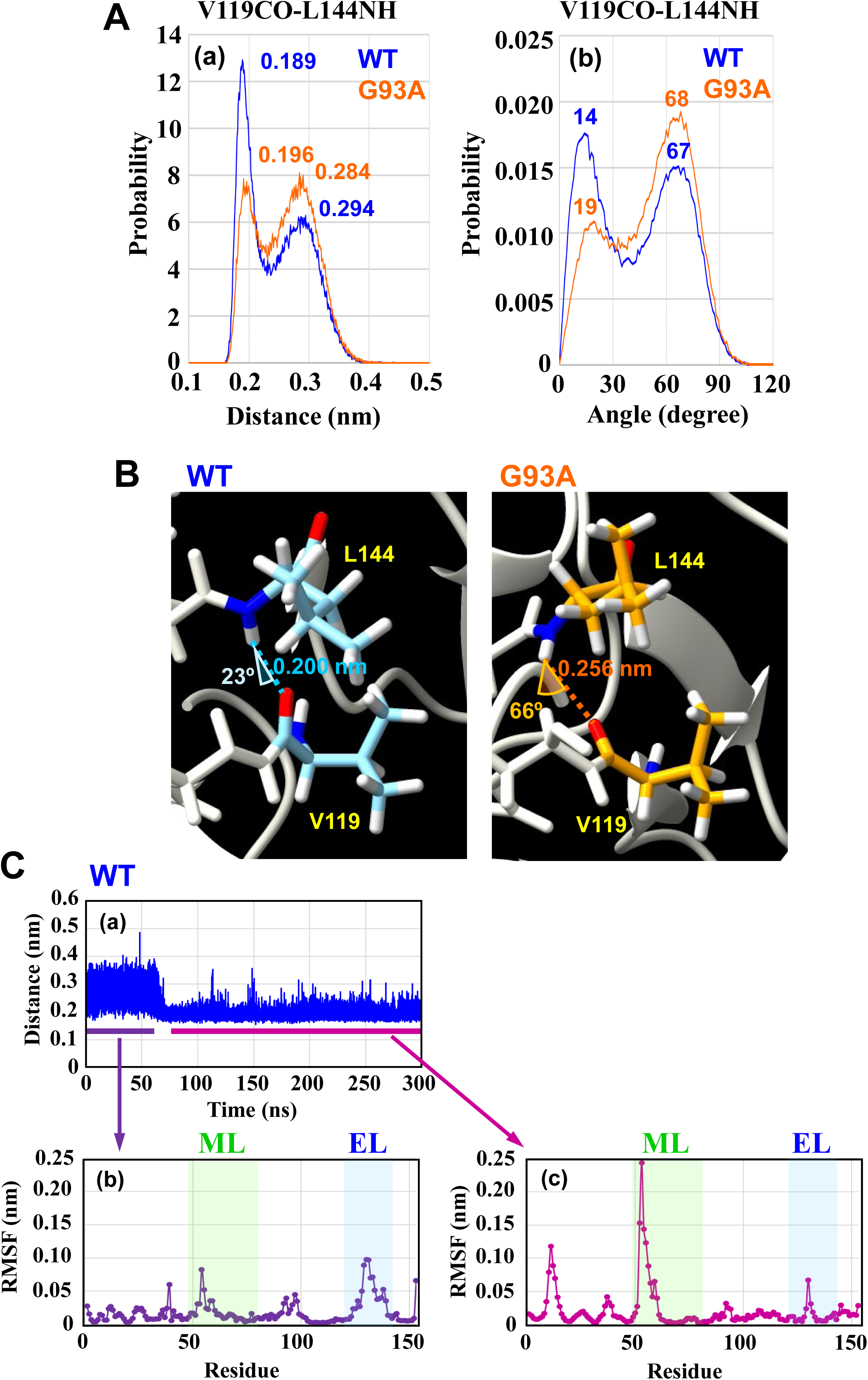
Change in Hydrogen-Bond Geometry between V119 and L144. ***A***, Hydrogen bonds regarding the C=O group of V119 and the N-H group of L144 in all geometries (*n*=150,000) from Run 1 to Run 5. (a), Distribution of the hydrogen-bond distances between the C=O group and the N-H group, (b), Distribution of the O-H-N angles of the C=O group and the N-H group. WT is drawn in blue and G93A is drawn in orange. The peak apex is indicated in nm. ***B***, Three-dimensional arrangement of the C=O group of V119 and the N-H group of L144 in WT and in G93A. The geometries are those of the cluster 1 of Run 1 in WT and G93A. ***C,*** (a), Time course of the hydrogen-bond distances between the C=O group of V119 and the N-H group of L144 in Run 1 of WT simulation. (b), RMSF (eigenvectors) of principal component 1 in Run 1 of WT simulation for 0 – 60 ns. (c), RMSF (eigenvectors) of principal component 1 in Run 1 of WT simulation for 70 – 300 ns.

Figure S7 shows the time course of the O^…^H distance between the C=O group of V119 and the NH group of L144, with hydrogen-bond formation typically indicated when the distance is around 0.2 nm. We classified the hydrogen-bond formation observed in these time courses into three types: instantaneous, short-term (lasting less than 50 ns), and long-term (lasting more than 150 ns). Given that the distance distribution for V119-L144 exhibits a characteristic time course, we compared the results of principal component analysis (PCA) for different patterns of hydrogen-bond formation (Figs. S8-1 to S8-10-7).

Figure 6C(a) illustrates the characteristic time course in the O^…^H distance in WT (corresponding to Run 1 in Fig. S7). Initially, the O^…^H distance fluctuates between 0.2 and 0.4 nm, which indicates that hydrogen-bond formation is only instantaneous. The result of PCA (Fig. S8-1-1) shows that no characteristic large fluctuations were observed in the interval from 0 to 60 ns (Fig. 6C(b)). However, after approximately 70 ns, the O^…^H distance decreases to the range of 0.2–0.3 nm and remains at this value until the end of the simulation, signifying long-term hydrogen-bond formation (Fig. 6C(a) and WT Run 1 in Fig. S7). At this period, substantial fluctuations occurred in residues 49–62 (a part of metal-binding loop) (Fig. 6C(c) and Fig. S8-1-2). This relationship between the hydrogen-bond time course and the PCA suggests that long-term hydrogen-bond formation between V119 and L144 is closely associated with large fluctuations in the metal-binding loop. A similar relationship was observed in WT Run 4 (Figs. S7 Run 4 in WT and S8-4-2) and in WT Run 5 (Figs S7 Run 5 in WT and S8-5-2).

In contrast, for G93A Runs 1, 2, 3 and 4, the O^…^H distance between the C=O group of V119 and the NH group of L144 fluctuated within 0.2 and 0.4 nm range with hydrogen-bond forming only instantaneously throughout these simulation periods, except after 270 ns in Run 3 (Figs. S7 Runs 1, 2, 3 and 4 in G93A). In Run 1 of G93A (Fig. S8-6), the fluctuation of the N-terminus was relatively large, and no other characteristic fluctuations were noted. In both Runs 2 and 3, large fluctuations of the electrostatic loop were observed (Figs. S8-7, S8-8), while in Run 4 the region of loop V containing A93 exhibited considerable fluctuation (Fig. S8-9). In Run 5, the distance decreased during the periods of 70–100 ns, 150–170 ns, and 240–280 ns (Fig. S7 Run 5 in G93A), suggesting formation of short-term hydrogen-bond. During these periods, the fluctuations of the metal-binding loop were relatively large, particularly in the 170–280 ns interval (Figs. S8-10, S8-10-5 and S8-10-6).

These results indicate that, in WT, a clear relationship exists between long-term V119-L144 hydrogen-bond formation and increased fluctuations of the metal-binding loop. In contrast, while such long-term V119-L144 hydrogen-bond was not observed in G93A, a correlation appears to exist between the short-term hydrogen-bond formation and increased fluctuations of the metal-binding loop.

#### 3. The steric change in L38 increases the fluctuation of β-strand 8 via I35 and L8

I35 is immediately adjacent to L38, and our data show that the distance between the C_α_ atoms of I35 and L38 is shorter in G93A than in WT (Fig. 5A(a)). Given that I35 is in direct contact with L8 (Fig. 1C), any change in the position of I35 is expected to influence L8. Moreover, as shown in Figure 7, since L8 interacts with β-strand 8, a positional shift of I35 transmitted via L8 could impact the mobility of β-strand 8. To investigate this possibility, we compared the distances between L8 and its adjacent amino acid residues in both WT and G93A (Figs. S9-1 to S9-3). In particular, Figures S9-2 and S9-3 show that the distance distribution between the H_γ_ atom of L8 and atoms of A145 or C146 in β-strand 8 exhibits two distinct peaks. In WT, the distal peak (indicating a longer distance) is more prevalent, whereas in G93A the proximal and distal peaks occur in comparable proportions. These results suggest that the side chain of L8 adopts two conformations at similar ratios in G93A, in contrast to WT where the distal conformation is favored. Importantly, when the H_γ_ atom of L8 is positioned close to β-strand 8 (corresponding to the proximal peak), the methyl groups at the δ-position are forced to adopt positions farther from β-strand 8. This reorientation likely increases the mobility of β-strand 8 (Fig. 7). Consequently, we expect that the mobility of β-strand 8 is higher in G93A than in WT (Fig. 2C).

**Figure 7.**
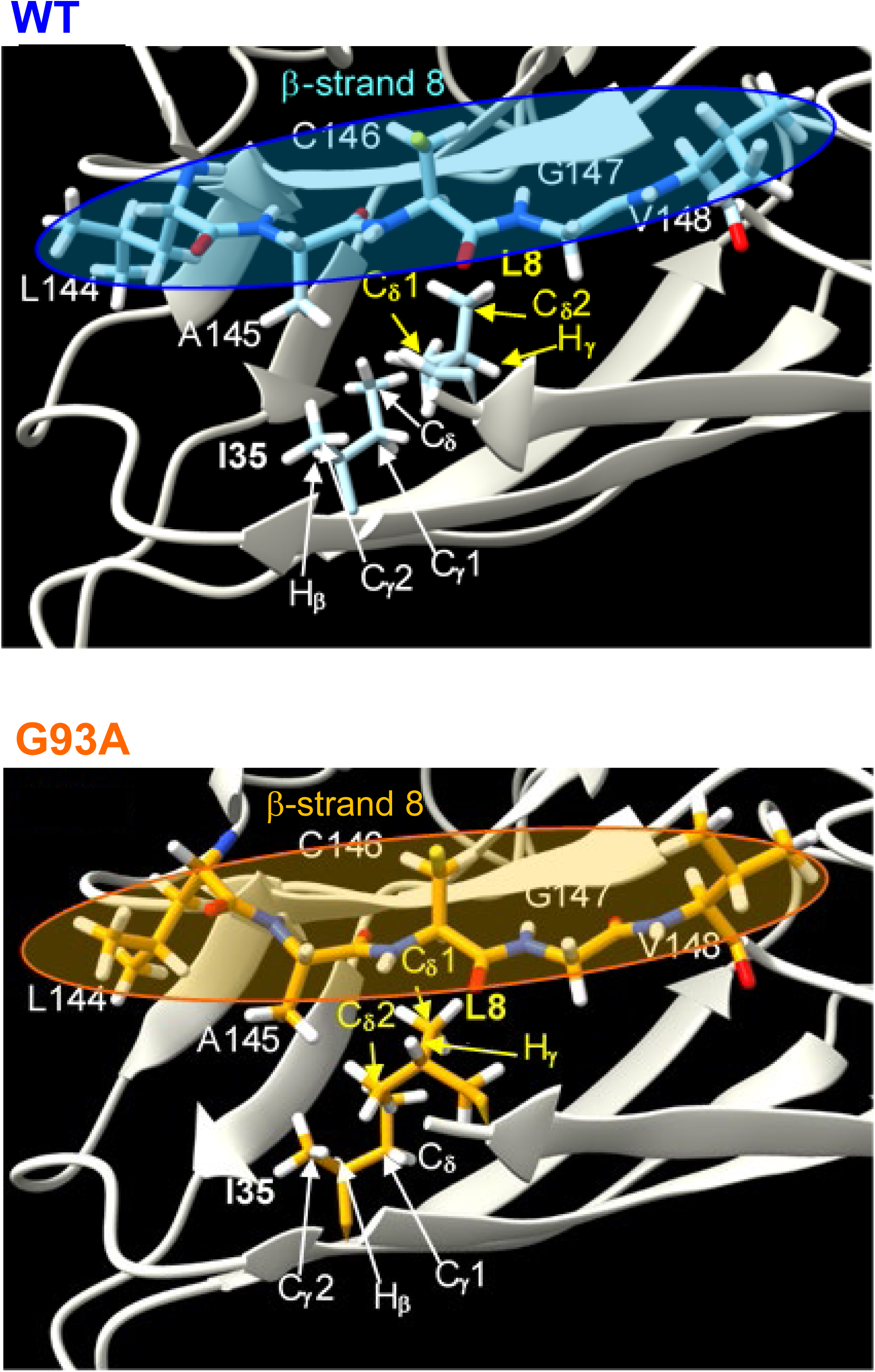
Comparison of Geometry of β-strand 8, I35, L8 in WT and G93A. Geometric arrangement of β-strand 8 (L144–V148) and hydrophobic residues I35 and L8 in Cluster 1 of Run 1 for WT (top) and G93A (bottom). The H_γ_, C_δ_1 and C_δ_2 atoms of L8 are highlighted in yellow.

### Introduction of the I35S mutation into G93A suppresses the fluctuation of the electrostatic loop

As we previously reported [22], the introduction of the I35S mutation into the G93A mutant suppressed aggregate formation in cells (Fig. S1). Furthermore, the eye degeneration observed in the G93A flies was suppressed by the additional I35S mutation (G93A/I35S) [24]. To understand these phenomena at the molecular level, we performed MD simulations and subsequent analyses on the G93A/I35S double mutant (Figs. S10-1 to S10-4). Figure 8A shows the positions of I35, L38 and L144 within the hydrophobic region in WT, G93A and G93A/I35S. Since the side chain of S35 is smaller than that of I35, which leads to several significant changes in the hydrophobic region. Firstly, the distance between the C_α_ of L38 and the C_α_ of S35 in G93A/I35S is 0.641 nm, which is shorter than the corresponding distance in G93A (0.739 nm) and WT (0.762 nm) (Fig. 8A). This reduction in the L38–I(S)35 distance is accompanied by a conformational change in the side chain of L38, attributable to a short distance between the C_α_ of L38 and the C_α_ of L144 in G93A/I35S (0.919 nm) compared with that in G93A (0.938 nm). In addition, this decreased distance between L38 and L144 alters the position of L144 and V119, thereby facilitating the formation of a long-term hydrogen bond between these residues, similar to that observed in WT in one of the five production runs for G93A/I35S (corresponding to Run 1 in Fig. S10-1). In this Run 1, significant fluctuations occurred in the electrostatic loop region during the 0–150 ns interval, when no hydrogen bond was formed between V119 and L144 (Fig. 8B(a), (b)). In contrast, as in the case of WT (Fig. 6C), when the hydrogen-bond was formed, substantial fluctuations were observed in residues 49–62 (a part of metal-binding loop) (Fig. 8B(c)). In fact, during the 170–300 ns interval in which the hydrogen bond was present, more pronounced fluctuations were observed across the entire metal-binding loop (Figs. S10-1 and S10-2, Run 1). Secondly, the RMSD of β-strand 8 in the G93A/I35S double mutant is larger than that of WT but smaller than that of G93A (Fig. 8C), indicating an intermediate level of fluctuation. The RMSD values for the electrostatic loop in G93A/I35S (Figs. 8D and S10-3) were lower than those for G93A (Fig. 2B). Furthermore, the difference between these values and the fluctuations observed in the metal-binding loop has diminished, which is a pattern similar to that observed in WT (Fig. 2B). This behavior can be attributed to the fact that replacing I35 with S35 increases the mobility of neighboring residues, while the formation of the hydrogen bond between V119 and L144 acts to enhance fluctuations of the metal-binding loop and suppress fluctuations of the electrostatic loop in the G93A/I35S mutant.

**Figure 8.**
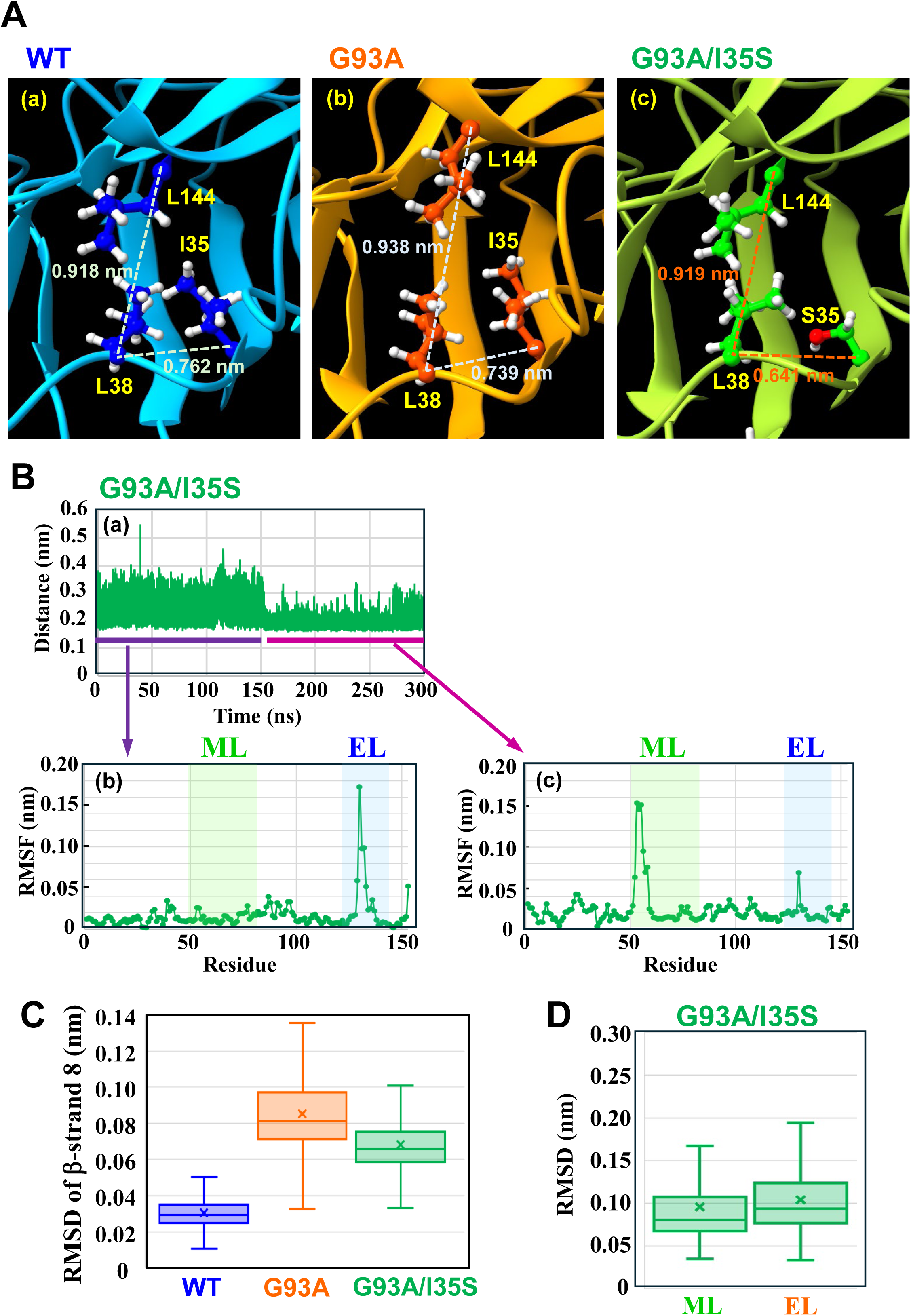
Comparison of Geometry of WT, G93A, and G93A/I35S. ***A*,** Geometry of Hydrophobic Region Including I35, L38, and L144. (a), Cluster 1 of Run 1 in WT simulation. (b), Cluster 1 of Run 1 in G93A simulation. (c), Cluster 1 of Run 1 in G93A/I35S simulation. ***B***, (a), Time course of the hydrogen-bond distances between the C=O group of V119 and the N-H group of L144 in Run 1 of G93A/I35S simulation. (b), RMSF (eigenvectors) of principal component 1 in Run 1 of G93A/I35S simulation for 0 – 150 ns. (c), RMSF (eigenvectors) of principal component 1 in Run 1 of G93A/I35S simulation for 160 – 300 ns. ***C***, Box-and-whisker plot of RMSD of the β-strand 8 in all geometries (*n*=150,000) from Run 1 to Run 5. The box represents IQR (interquartile range: 25-75th percentile), the horizontal line inside the box represents the median value, and the cross mark represents the average value. The top of the whiskers above the box represents the maximum value, and that below the bottom represents the minimum value. WT, G93A, and G93A/I35S are drawn in blue, orange, and green, respectively. ***D***, Box-and-whisker plot of RMSD of the metal-binding loop (ML: 49-82) and the electrostatic loop (EL: 121-142) of G93A/I35S mutant in all geometries (*n*=150,000) from Run 1 to Run 5. The box represents IQR (interquartile range: 25-75th percentile), the horizontal line inside the box represents the median value, and the cross mark represents the average value. The top of the whiskers above the box represents the maximum value, and that below the bottom represents the minimum value.

## Discussion

In this study, we compared the structures of WT, G93A, and G93A/I35S SOD1 monomers using MD simulations. The results demonstrate that the single-point mutation from G93 to A93 alters the fluctuations of both the electrostatic loop and the metal-binding loop by inducing conformational changes in the hydrophobic region surrounding L38. Figure 9 shows schematic representations of the loop shapes, characteristic fluctuations, and hydrophobic regions in WT, G93A, and G93A/I35S. Replacing G93 with A93 leads to a repositioning of the adjacent L38 side chain. This positional change in L38, in turn, affects nearby residues such as I35, V87, and L144 (Figs. 5, 9(b) and 9(d)), thereby modifying their spatial arrangement. When the position of I35 is altered, the mobility of the L8 side chain increases (Fig. S9), which subsequently enhances the fluctuation of β-strand 8 (Fig. 2C). Since β-strand 8 is directly connected to the electrostatic loop (Fig. 1C), its increased mobility can be transmitted to, and thereby amplify, the fluctuations of the electrostatic loop (Figs. 2B and 3).

**Figure 9.**
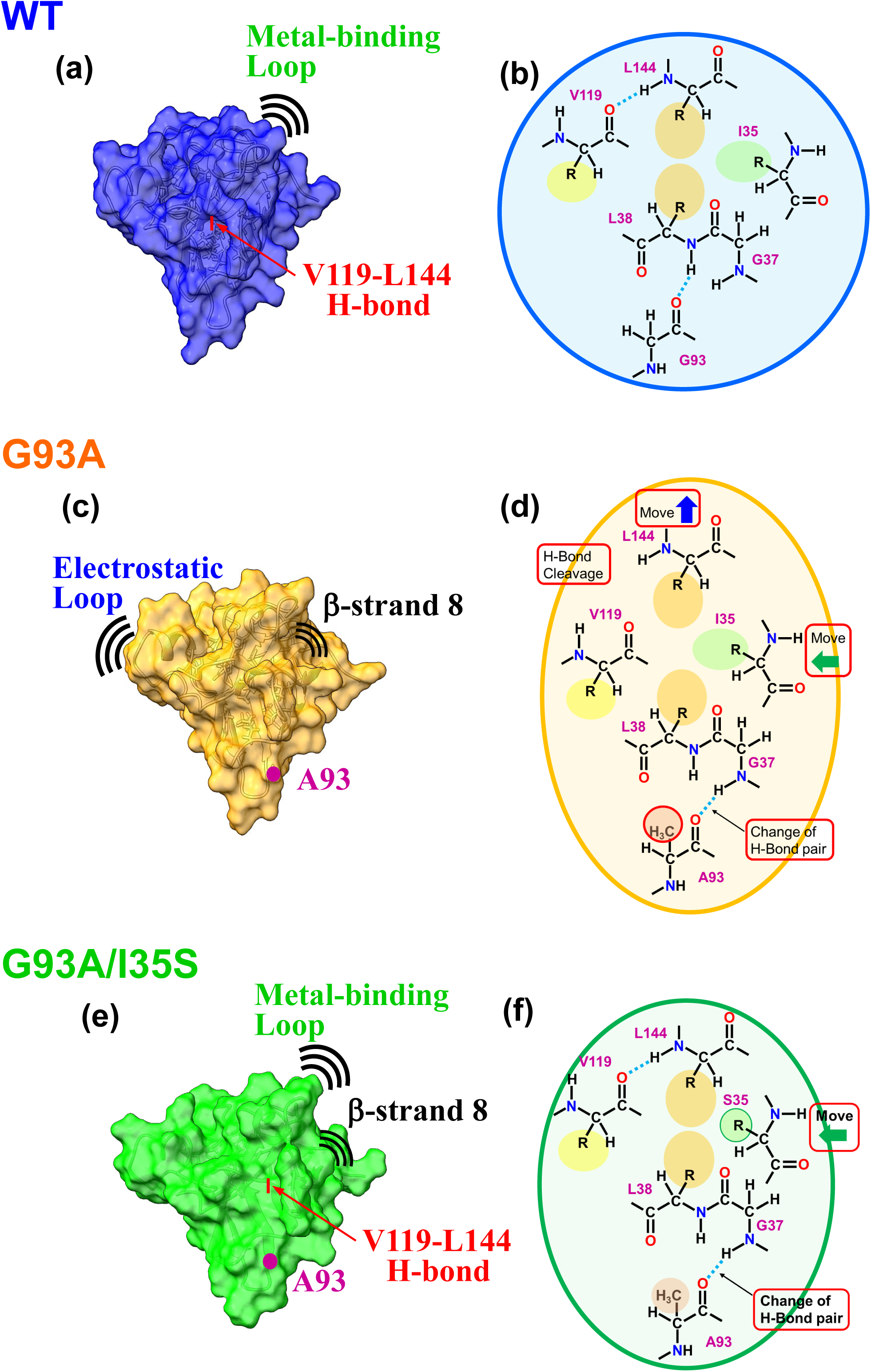
Summary of Differences in Hydrophobic Region of WT, G93A, and G93A/I35S. Geometry and characteristic fluctuation of the loop and schematic representation of hydrophobic region in WT (a), (b), in G93A (c), (d), and in G93A/I35S (e), (f).

Furthermore, the change in the position of L38 also induces a shift in the position of L144, which critically affects the formation of a stabilizing hydrogen bond between V119 and L144 (Fig. 6). In WT SOD1, the orientation of the NH group of L144 relative to the C=O group of V119 is favorable for hydrogen-bond formation, resulting in long-term bond formation (Fig. 6C). The hydrogen bond, located between β-strand 8 and the electrostatic loop, is thought to act as a buffer that suppresses the transmission of excessive fluctuations from β-strand 8 to the electrostatic loop. In G93A, however, the altered position of L144 makes the formation of the V119-L144 hydrogen bond less favorable, leading to more pronounced transmission of fluctuations from β-strand 8 to the electrostatic loop. This mechanism is supported by our PCA results (Fig. S8), which indicate that when the overall fluctuations are concentrated in one loop, the fluctuations in the other loop tend to be reduced.

When I35 is mutated to S35 in the G93A background (G93A/I35S), the distance between L38 and S35 becomes shorter than the corresponding distances observed in both WT and G93A (Fig. 8A). This reduction in distance decreases the steric crowding around L8, resulting in large fluctuations of β-strand 8 similar to those observed in G93A (Fig. 8C). However, in the G93A/I35S double mutant, the distance between L38 and L144 is restored to a value comparable to that of WT (Figs. 8A, 9(b) and 9(f)), and the formation of the hydrogen bond between V119 and L144 is enhanced compared to G93A. This restoration of the V119-L144 hydrogen bond reduces the fluctuation of the electrostatic loop and enhances the metal-binding loop (Fig. 8B), even though β-strand 8 remains relatively mobile, resulting in a fluctuation pattern that resembles that of WT (Figs. 2B and 8D). Given that increased fluctuations of the electrostatic loop have been linked to SOD1 aggregation [14], our findings provide a molecular mechanism explaining the experimental observation that aggregation in G93A is suppressed in the G93A/I35S double mutant [22].

According to Souza *et al*., the residues T39, H43, K122, A123, A140 and G141 are involved in the network that explains the action of the distant G93A mutation on the electrostatic loop [29]. In our analysis, H43 engages in two hydrogen bonds with V87 (Figs. S11-1 and S11-2) and also tends to form stable hydrogen bonds with H120. Furthermore, G44, an adjacent residue to H43, forms stable backbone hydrogen bonds with K122 and H120. In addition, H120, E121, and K122 establish stable hydrogen bonds with G141, S142, and A140, respectively; residues that constitute the other chain of the electrostatic loop. This network suggests that changes in the position of V87 (which neighbors L38 and is altered by the G93A mutation; see Fig. 5A(b) and Fig. 5B) are likely transmitted to the electrostatic loop via H43 and G44. Therefore, if the conformational change of H43 alters the geometry of the electrostatic loop, then the altered position of V87 is expected to have a more pronounced effect than changes in T39.

Our MD simulations indicate that the conformational changes in the hydrophobic region, centered on L38, induced by the G93A mutation (Fig. 4 and 5) directly affect the fluctuation of the electrostatic loop. To further validate these findings, we compared the structural changes observed in the simulations with the X-ray crystal structures of WT SOD1 (PDB ID: 3T5W) [45] and G93A (PDB ID: 3GZO) [46] (Fig. S12). Table S12 demonstrates that the shorter distance between the C=O group of G93 (or A93) and the N atom of L38 in WT, as well as the shorter distance between the C=O group of G93 (or A93) and the N atom of G37 in G93A, support the altered hydrogen-bonding patterns shown in Figure 4. Although the differences in the L38-L144 distance were comparable to the simulation results, the distances between L38-I35 and L38-V87 were both shorter in G93A than in WT, which is consistent with the hydrophobic region alterations observed in our simulations. These analyses indicate that, despite minor differences in structural features, there is reasonable agreement between the simulation results and the X-ray structures.

Numerous studies have demonstrated that the misfolding and aggregation of both mutant SOD1 and even WT SOD1 are key processes in ALS pathogenesis [47–50]. Aggregation of SOD1 may occur through several mechanisms, including aberrant hydrogen-bond formation [51, 52], oxidative modification [53, 54], and cleavage of the intramolecular disulfide S-S bond (C57–C146) [21, 55, 56]. Recent cryo-EM studies have revealed the atomic-level structure of amyloid fibrils formed by WT SOD1 [56]. These analyses showed that the conformations of the electrostatic and metal-binding loops are completely altered and that intermolecular interactions, including hydrogen bonds, drive the formation of SOD1 fibrils. However, the process by which oligomeric species transform into amyloid fibrils remains incompletely understood. Our observation that the electrostatic loop in G93A exhibits increased fluctuations suggests a mechanism whereby these dynamic changes may lead to metal ion loss from mature SOD1, thereby facilitating oligomerization through enhanced intermolecular interactions.

This study further revealed that small structural changes in the hydrophobic region, including alterations in L38, I35, and L144, lead to increased fluctuations of both the electrostatic loop and β-strand 8 in G93A. Moreover, our previous work has shown that I35 plays an essential role in intracellular aggregation of ALS mutant SOD1 [22]. In support of this, Srinivasan *et al*. reported that the small-molecule compound, resveratrol, reverses structural changes in G93A [57]. Their docking studies, based on MD simulations, indicated that resveratrol preferentially binds to the exterior of the β-sheet rather than the loop of SOD1. Since the interior of the β-sheets forms a hydrophobic region, these findings suggest that resveratrol may induce a conformational change in this region, which is potentially related to the recovery of normal SOD1 structure. Therefore, targeting the hydrophobic region, including residue I35, represents a promising strategy for ALS therapeutic development.

### Concluding remarks

In this study, we revealed distinct fluctuation patterns in the electrostatic and metal-binding loops of SOD1 monomers for WT, G93A, and G93A/I35S based on five independent 300-ns atomistic MD simulations. Characteristic fluctuations were observed in the metal-binding loop in both WT and G93A/I35S double mutant, whereas in G93A the electrostatic loop displayed pronounced fluctuations. Our analyses indicate that a seemingly minor substitution from G93 to A93 results in large differences in activation of electrostatic loop fluctuations *via* alterations in hydrophobic interactions. Moreover, the formation, or disruption, of the V119-L144 hydrogen bond plays a pivotal role in switching the fluctuation pattern between these loops. The behavior of the G93A/I35S double mutant, specifically the suppression of aggregate formation observed in our previous report [22], might be explained by the restoration of the structural integrity of the hydrophobic region and the reformation of the V119-L144 hydrogen bond. Notably, the large structural changes observed in the electrostatic loop of fALS mutant SOD1 are likely to promote aggregate formation through enhanced intermolecular association. Overall, these findings further our understanding of how structural changes in fALS mutant SOD1 contribute to the pathogenesis of ALS.

### Methods: Details of Molecular Dynamics Simulation

The starting structure for the WT SOD1 monomer was obtained from a 1.8 Å crystal structure of the chain A of human SOD1 (PDB ID: 3T5W) [45]. The ionization states of the amino acids corresponded to neutral pH (Asp^−^, Glu^−^, His^0^, Lys^+^, and Arg^+^). The C57-C146 disulfide bond was intact, consistent with in vitro studies. The modified CME 111 residue in the PDB structure was reverted to the unmodified Cys. Graphics of SOD1 and its mutants were generated using UCSF ChimeraX 1.5 [58, 59].

Settings of the initial system were made using the Amber Tools19 package [60]. The AMBER14SB force field was selected for the simulations of the protein [61]. Metal-ligand interaction parameters for Zn^2+^ and Cu^+^ ions were generated using the MCPB.py program [62]. The protein molecules were solvated in a rectangular box with TIP3P water molecules [63], and the system was neutralized by adding Na^+^ ions using the tLEaP program.

Molecular dynamics (MD) simulations of the wild-type SOD1 protein and its mutants were performed using the GROMACS 2019 package [64]. The energy of the initial system was minimized using the steepest descent method. After minimizing the energy of the system, the system was equilibrated temperature at 310 K using the V-rescale thermostat [65] during 100 ps (*NVT* equilibrium) and then equilibrated pressure at 1.0 bar using the Parrinello-Rahman barostat [66] during 100 ps (*NPT* equilibrium). The production runs were further performed at a temperature of 310 K with the V-rescale thermostat and the Parrinello-Rahman barostat for 300 ns using the LINCS (linear constraint solver) [67] for constraint of X-H bond distances and PME (particle mesh Ewald) [68] algorithms for calculation of Coulomb energy, and the position and the velocity were recorded every 100 ps. The production runs were performed five times for each system: WT, G93A, and the G93A/I35S double mutant.

The obtained MD trajectories were analyzed using the following GROMACS distribution programs: gmx rms, gmx rmsf, gmx cluster, gmx covar, gmx anaeig, and gmx distance. These MD analyses generated parameter values for the root-mean-square deviation (RMSD) and the root-mean-square fluctuation (RMSF). Cluster analyses were performed using the RMSD-based Gromos method with a cutoff of 0.100 nm [69]. Interatomic distances were analyzed by plotting the distributions at 30001 points obtained from five production runs.

## Abbreviations

MD: molecular dynamics
SOD: superoxide dismutase
ALS: amyotrophic lateral sclerosis
RMSD: root-mean-square deviation
RMSF: root-mean-square fluctuation
PCA: principal component analysis

## References

1. McCord, J. M. and Fridovich, I. (1969) Superoxide Dismutase, J. Biol. Chem. 244, 6049–6055

2. Tainer, J. A., Getzoff, E. D., Richardson, J. S., and Richardson, D. C. (1983) Structure and mechanism of copper, zinc superoxide dismutase, Nature 306, 284–287

3. Rosen, D. R., Siddique, T., Patterson, D., Figlewicz, D. A., Sapp, P., Hentati, A., Donaldson, D., Goto, J., O’Regan, J. P., Deng, H.-X., Rahmani, Z., Krizus, A., McKenna-Yasek, D., Cayabyab, A., Gaston, S. M., Berger, R., Tanzi, R. E., Halperin, J. J., Herzfeldt, B., Bergh, R. V. d., Hung, W.-Y., Bird, T., Deng, G., Mulder, D. W., Smyth, C., Laing, N. G., Soriano, E., Pericak-Vance, M. A., Haines, J., Rouleau, G. A., Gusella, J. S., Horvitz, H. R., and Bound Jr, R. H. (1993) Mutations in Cu/Zn superoxide dismutase gene are associated with familial amyotrophic lateral sclerosis, Nature 362, 59–62

4. Shaw, B. F., Lelie, H. L., Durazo, A., Nersissian, A. M., Xu, G., Chan, P. K., Gralla, E. B., Tiwari, A., Hayward, L. J., Borchelt, D. R., Valentine, J. S., and Whitelegge, J. P. (2008) Detergent-insoluble Aggregates Associated with Amyotrophic Lateral Sclerosis in Transgenic Mice Contain Primarily Full-length, Unmodified Superoxide Dismutase-1, J. Biol. Chem. **283**, 8340–8350

5. Wright, G. S. A., Antonyuk, S. V., and Hasnain, S. S. (2019) The biophysics of superoxide dismutase-1 and amyotrophic lateral sclerosis, Quart. Rev. Biophys. 52, e12, 1–39

6. Reaume, A. G., Elliott, J. L., Hoffman, E. K., Kowall, N. W., Ferrante, R. J., Siwek, D. F., Wilcox, H. M., Flood, D. G., Beal, M. F., Brown Jr., R. H., Scott, R. W., and Snider, W. D. (1996) Motor neurons in Cu/Zn superoxide dismutase-deficient mice develop normally but exhibit enhanced cell death after axonal injury. Nat Genet., 13, 43–47

7. Richardson, J. S., Thomas, K. A., Rubin, B. H., and Richardson, D. C. (1975) Crystal Structure of Bovine Cu,Zn Superoxide Dismutase at 3 Å Resolution: Chain Tracing and Metal Ligands, Proc. Nat. Acad. Sci. U. S. A. **72**, 1349–1353

8. Valentine, J. S., Doucette, P. A., and Potter, S. Z. (2005) Copper-Zinc Superoxide Dismutase and Amyotrophic Lateral Sclerosis, Annu. Rev. Biochem. 74, 563–593

9. Tainer, J. A., Getzoff, E. D., Beem, K. M., Richardson, J. S., and Richardson, D. C. (1982) Determination and Analysis of the 2 Å Structure of Copper, Zinc Superoxide Dismutase, J. Mol. Biol. 160, 181–217

10. Getzoff, E. D., Tainer, J. A., Weiner, P. K., Kollman, P. A., Richardson, J. S., and Richardson, D. C. (1983) Electrostatic recognition between superoxide and copper, zinc superoxide dismutase, Nature 306, 287–290

11. Strange, R. W., Antonyuk, S., Hough, M. A., Doucette, P. A., Rodriguez, J. A., Hart, P. J., Hayward, L. J., Valentine, J. S., and Hasnain, S. S. (2003) The Structure of Holo and Metal-deficient Wild-type Human Cu, Zn Superoxide Dismutase and Its Relevance to Familial Amyotrophic Lateral Sclerosis, J. Mol. Biol. 328, 877–891

12. Johnston, J. A., Dalton, M. J., Gurney, M. E., and Kopito, R. R. (2000) Formation of high molecular weight complexes of mutant Cu,Zn-superoxide dismutase in a mouse model for familial amyotrophic lateral sclerosis, Proc. Natl. Acad. Sci. U. S. A. 97, 12571–12576

13. Berdyński, M., Miszta, P., Safranow, K., Andersen, P. M., Morita, M., Filipek, S., Żekanowski, C., and Kuźma-Kozakiewicz, M., (2022) SOD1 mutations associated with amyotrophic lateral sclerosis analysis of variant severity, Scientific Reports 12, 103

14. Molnar, K. S., Karabacak, N. M., Johnson, J. L., Wang, Q., Tiwari, A., Hayward, L. J., Coales, S. J., Hamuro, Y., and Agar, J. N. (2009) A Common Property of Amyotrophic Lateral Sclerosis-associated Variants Destabilization of the Copper/Zinc Superoxide Dismutase Electrostatic Loop, J. Biol. Chem. 284, 30965–30973

15. Crow, J. P., Sampson, J. B., Zhuang, Y., Thompson, J. A., and Beckman, J. S. (1997) Decreased Zinc Affinity of Amyotrophic Lateral Sclerosis-Associated Superoxide Dismutase Mutants Leads to Enhanced Catalysis of Tyrosine Nitration by Peroxynitrite, J. Neurochem. 69, 1936–1944

16. Mulligan, V. K., Kerman, A., Ho, S., and Chakrabartty, A. (2008) Denaturational Stress Induces Formation of Zinc-Deficient Monomers of Cu,Zn Superoxide Dismutase: Implications for Pathogenesis in Amyotrophic Lateral Sclerosis, J. Mol. Biol. 383, 424–436.

17. Khare, S. D., Caplow, M., and Dokholyan, N. V. (2004) The rate and equilibrium constants for a multistep reaction sequence for the aggregation of superoxide dismutase in amyotrophic lateral sclerosis, Proc. Natl. Acad. Sci. U. S. A. 101, 15094–15099

18. Arnesano, F., Banci, L., Bertini, I., Martinelli, M., Furukawa Y., and, O’Halloran T. V. (2004) The Unusually Stable Quaternary Structure of Human Cu,Zn-Superoxide Dismutase 1 Is Controlled by Both Metal Occupancy and Disulfide Status, J. Biol. Chem. **279**, 47998–48003

19. Lindberg, M. J., Normark, J., Holmgren, A., and Oliveberg, M. (2004) Folding of human superoxide dismutase: Disulfide reduction prevents dimerization and produces marginally stable monomers, Proc. Natl. Acad. Sci. U. S. A. 101, 15893–15898

20. Sheng, Y., Chattopadhyay, M., Whitelegge, J., and Valentine, J. S. (2012) SOD1 Aggregation and ALS: Role of Metallation States and Disulfide Status, Curr. Top. Med. Chem. 12, 2560–2572

21. Fujiwara, N., Wagatsuma, M., Oba, N., Yoshihara, D., Tokuda, E., Sakiyama, H., Eguchi, H., Ichihashi, M., Furukawa, Y., Inoue, T., and Suzuki, K. (2018) Cu/Zn-superoxide dismutase forms fibrillar hydrogels in a pH-dependent manner via a water-rich extended intermediate state, PLoS One, 13, e0205090.

22. Asai, Y., Yano, K., Higashino, T., Yoshihara, D., Sakiyama, H., Eguchi, H., Fukushima, K., Suzuki, K., and Fujiwara, N. (2025) The Ile35 residue of the ALS-associated mutant SOD1 plays a crucial role in the intracellular aggregation of the molecule, Mol. Neurobiol. 62, 2023–2038. doi: 10.1007/s12035-024-04369-0.

23. McAlary L, Nan JR, Shyu C, Sher M, Plotkin SS, Cashman NR. (2024) Amyloidogenic regions in beta-strands II and III modulate the aggregation and toxicity of SOD1 in living cells. Open Biol. Jun;14(6):230418. doi: 10.1098/rsob.230418.

24. Yano K, Fujino Y, Nagai Y, Fujiwara, N. (2026) Ile35 residue of SOD1 plays a key role in aggregate formation and eye degeneration in a Drosophila model of ALS. *BioRxiv* doi: 10.64898/2026.09.13.751333

25. Karplus, M. and Petsko, G. A. (1990) Molecular dynamics simulations in biology, Nature 347, 631−639

26. Karplus, M. and McCammon, J. A. (2002) Molecular dynamics simulations of biomolecules, Nature Struct.Biol. 9, 646−651

27. Adcock, S. A. and McCammon, J. A. (2006) Molecular Dynamics: Survey of Methods for Simulating the Activity of Proteins, Chem. Rev. 106, 1589−1615

28. Hollingsworth, S. A. and Dror, R. O. (2018) Molecular Dynamics Simulation for All, Neuron 99, 1129−1143

29. Souza, P. C. T., Thallmair, S., Marrink, S. J., and Mera-Adasme, R. (2019) An Allosteric Pathway in Copper, Zinc Superoxide Dismutase Unravels the Molecular Mechanism of the G93A Amyotrophic Lateral Sclerosis-Linked Mutation, J. Phys. Chem. Lett. **10**, 7740−7744

30. Khare, S. D. and Dokholyan, N. V. (2006) Common dynamical signatures of familial amyotrophic lateral sclerosis-associated structurally diverse Cu, Zn superoxide dismutase mutants, Proc. Natl. Acad. Sci. U. S. A. 103, 3147–3152

31. Strange, R. W., Yong, C. W., Smith, W., and Hasnain, S. S. (2007) Molecular dynamics using atomic-resolution structure reveal structural fluctuations that may lead to polymerization of human Cu–Zn superoxide dismutase, Proc. Natl. Acad. Sci. U. S. A. 104, 10040–10044

32. Ding, F. and Dokholyan, N. V. (2008) Dynamical roles of metal ions and the disulfide bond in Cu, Zn superoxide dismutase folding and aggregation, Proc. Natl. Acad. Sci. U. S. A. 105, 19696 –19701

33. Schmidlin, T., Kennedy, B. K., and Daggett, V. (2009) Structural Changes to Monomeric CuZn Superoxide Dismutase Caused by the Familial Amyotrophic Lateral Sclerosis-Associated Mutation A4V, Biophys. J. 97, 1709–1718

34. Nedd, S., Redler, R. L., Proctor, E. A., Dokholyan, N. V., and Alexandrova, A. N. (2014) Cu,Zn-Superoxide Dismutase without Zn Is Folded but Catalytically Inactive, J. Mol. Biol. 426, 4112–4124

35. Mera-Adasme, R., Erdmann, H., Bereźniak, T., and Ochsenfeld, C. (2016) Destabilization of the metal site as a hub for the pathogenic mechanism of five ALS-linked mutants of copper, zinc superoxide dismutase, Metallomics 8, 1141–1150

36. Healy, E. F. (2017) A prion-like mechanism for the propagated misfolding of SOD1 from in silico modeling of solvated near-native conformers, PLoS ONE 12, e0177284

37. Pereira, G. R. C., Tellini, G. H. A. S., and De Mesquita, J. F. (2019) *In silico* analysis of PFN1 related to amyotrophic lateral sclerosis, PLoS ONE 14, e0215723

38. Srinivasan, E. and Rajasekaran, R. (2019) Computational Investigation on Electrostatic Loop Mutants Instigating Destabilization and Aggregation on Human SOD1 Protein Causing Amyotrophic Lateral Sclerosis, Protein J. 38, 37–49

39. Jahan, I. and Nayeem, S. M. (2020) Conformational dynamics of superoxide dismutase (SOD1) in osmolytes: a molecular dynamics simulation study, RSC Adv. 10, 27598–27614

40. Pereira, G. R. C., Vieira, B. de A. A., and De Mesquita, J. F. (2021) *Comprehensive in silico* analysis and molecular dynamics of the superoxide dismutase 1 (SOD1) variants related to amyotrophic lateral sclerosis, PLoS ONE 16, e0247841

41. Basith, S., Manavalan, B., and Lee, G. (2022) Amyotrophic lateral sclerosis disease-related mutations disrupt the dimerization of superoxide dismutase 1 – A comparative molecular dynamics simulation study, Comput. Biol. Med. 151, 106319

42. Alemasov, N. A., Timofeev, V. S., Ivanisenko, N. V., Kolchanov, N. A., and Ivanisenko, V. A. (2022) Computer analysis of the relation between hydrogen bond stability in SOD1 mutants and the survival time of amyotrophic lateral sclerosis patients, J. Mol. Graph. Model. 110, 108026

43. Kalia, M., Miotto, M., Ness, D., Opie-Martin, S., Spargo, T. P., Rienzo, L. D., Biagini, T., Petrizzelli, F., Al Khleifat, A., Kabiljo, R., Project MinE ALS Sequencing Consortium, SOD-ALS clinical and genetic data collection group, Mazza, T., Ruocco, G., Milanetti E., Dobson, R. J. B., Al-Chalabi, A., Lacoangeli, A. (2023) Molecular dynamics analysis of superoxide dismutase 1 mutations suggests decoupling between mechanisms underlying ALS onset and progression, Comput. Struct. Biotech. J. **21**, 5296-5308

44. Zhu, J., Wang, J., Han, W., Xu, D. (2022) Neural relational inference to learn long-range allosteric interactions in proteins from molecular dynamics simulations, Nature Commun. 13, 1661

45. Ihara, K., Fujiwara, N., Yamaguchi, Y., Torigoe, H., Wakatsuki, S., Taniguchi, N., and Suzuki, K. (2012) Structural switching of Cu,Zn-superoxide dismutases at loop VI: insights from the crystal structure of 2-mercaptoethanol-modified enzyme, Biosci. Rep. **32**, 539–548

46. Galaleldeen, A., Strange, R. W., Whitson, L. J., Antonyuk, S. V., Narayana, N., Taylor, A. B., Schuermann, J. P., Holloway, S. P., Hasnain, S. S., Hart, P. J. (2009) Structural and biophysical properties of metal-free pathogenic SOD1 mutants A4V and G93A, Arch. Biochem. Biophys. 492, 40–47

47. Julien, JP. (2001) Amyotrophic lateral sclerosis: Unfolding the toxicity of the misfolded. Cell. 104, 581–591

48. Tokuda, E., Takei, Yi., Ohara, S., Fujiwara, N., Furukawa, Y. (2019) Wild-type Cu/Zn-superoxide dismutase is misfolded in cerebrospinal fluid of sporadic amyotrophic lateral sclerosis. Mol Neurodegeneration. 14, 42

49. Furukawa, Y., Kaneko, K., Yamanaka, K., Nukina, N. (2010) Mutation-dependent polymorphism of Cu,Zn-superoxide dismutase aggregates in the familial form of amyotrophic lateral sclerosis. J. Biol. Chem. 285, 22221–22231.

50. Nagano, S., Takahashi, Y., Yamamoto, K., Masutani, H., Fujiwara, N., Urushitani, M., Araki, T. (2015) A cysteine residue affects the conformational state and neuronal toxicity of mutant SOD1 in mice: Relevance to the pathogenesis of ALS. Hum. Mol. Genet. 24, 3427–3439.

51. Banci, L., Bertini, I., Durazo, A., Girotto, S., Gralla, E. B., Martinelli, M., Valentine, J. S., Vieru, M., Whitelegge, J. P. (2007) Metal-free superoxide dismutase forms soluble oligomers under physiological conditions: A possible general mechanism for familial ALS, Proc. Natl. Acad. Sci. U. S. A. 104, 11263–11267

52. Elam, J. S., Taylor, A. B., Strange, R., Antonyuk, S., Doucette, P. A., Rodriguez, J. A., Hasnain, S. S., Hayward, L. J., Valentine, J. S., Yeates, T. O., Hart, P. J. (2003) Amyloid-like filaments and water-filled nanotubes formed by SOD1 mutant proteins linked to familial ALS, *Nature Struct*. Biol. 10, 461–467

53. Fujiwara, N, Nakano, M, Kato, S, Yoshihara, D, Ookawara, T, Eguchi, H, Taniguchi, N, Suzuki, K. (2007) Oxidative modification to cysteine sulfonic acid of Cys111 in human copper-zinc superoxide dismutase. J Biol Chem. 282, 35933–35944.

54. Bosco DA, Morfini G, Karabacak NM, Song Y, Gros-Louis F, Pasinelli P, Goolsby H, Fontaine BA, Lemay N, McKenna-Yasek D, Frosch MP, Agar JN, Julien JP, Brady ST, Brown RH Jr. (2010) Wild-type and mutant SOD1 share an aberrant conformation and a common pathogenic pathway in ALS. Nat Neurosci. 13, 1396–403.

55. Furukawa, Y. (2021) Good and Bad of Cu/Zn-Superoxide Dismutase Controlled by Metal Ions and Disulfide Bonds, Chem. Lett. 50, 331–341

56. Wang, L.-Q., Ma, Y., Yuan, H.-Y., Zhao, K., Zhang, M.-Y., Wang, Q., Huang, X., Xu, W.-C., Dai, B., Chen, J., Li, D., Zhang, D., Wang, Z., Zou, L., Yin, P., Liu, C., and Liang, Y. (2022) Cryo-EM structure of an amyloid fibril formed by full-length human SOD1 reveals its conformational conversion, Nature Commun. 13, 3491

57. Srinivasan, E. and Rajasekaran, R. (2018) Quantum chemical and molecular mechanics studies on the assessment of interactions between resveratrol and mutant SOD1 (G93A) protein, J. Comput. Aided. Mol. Des. 32, 1347–1361

58. Goddard, T. D., Huang, C. C., Meng, E. C., Pettersen, E. F., Couch, G. S., Morris, J. H., and Ferrin, T. E. (2018) UCSF ChimeraX: Meeting modern challenges in visualization and analysis, Pro. Sci. 27, 14–25

59. Pettersen, E. F., Goddard, T. D., Huang, C. C., Meng, E. C., Couch, G. S., Croll, T. I., Morris, J. H., and Ferrin, T. E. (2021) UCSF ChimeraX: Structure visualization for researchers, educators, and developers, Pro. Sci. 30, 70–82

60. Case, D.A., Ben-Shalom, I.Y., Brozell, S.R., Cerutti, D.S., Cheatham, III, T.E., Cruzeiro, V.W.D., Darden, T.A., Duke, R.E., Ghoreishi, D., Giambasu, G., Giese, T., Gilson, M.K., Gohlke, H., Goetz, A.W., Greene, D., Harris, R., Homeyer, N., Huang, Y., Izadi, S., Kovalenko, A., Krasny, R., Kurtzman, T., Lee, T.S., LeGrand, S., Li, P., Lin, C., Liu, J., Luchko, T., Luo, R., Man, V., Mermelstein, D.J., Merz, K.M., Miao, Y., Monard, G., Nguyen, C., Nguyen, H., Onufriev, A., Pan, F., Qi, R., Roe, D.R., Roitberg, A., Sagui, C., Schott-Verdugo, S., Shen, J., Simmerling, C.L., Smith, J., Swails, J., Walker, R.C., Wang, J., Wei, H., Wilson, L., Wolf, R.M., Wu, X., Xiao, L., Xiong, Y., York, D.M., and Kollman, P.A. (2019) AMBER 2019, University of California, San Francisco.

61. Maier, J. A., Martinez, C., Kasavajhala, K., Wickstrom, L., Hauser, K.E., and Simmerling, C. (2015) ff14SB: Improving the Accuracy of Protein Side Chain and Backbone Parameters from ff99SB. J. Chem. Theory Comput. 11, 3696–3713

62. Li, P. and Merz, Jr., K. M. (2016) MCPB.py: A Python Based Metal Center Parameter Builder, J. Chem. Inf. Model. 56, 599–604

63. Jorgensen, W. L., Chandrasekhar, J., Madura, J. D., Impey, R. W., and Klein, M. L. (1983) Comparison of simple potential functions for simulating liquid water, J. Chem. Phys. 79, 926–935

64. Van Der Spoel, D., Lindahl, E., Hess, B., Groenhof, G., Mark, A. E., and Berendsen, H. J. C. (2005) GROMACS: Fast, Flexible, and Free, J. Comput. Chem. 26, 1701–1718

65. Bussi, G., Donadio, D., and Parrinello, M. (2007) Canonical sampling through velocity rescaling, J. Chem. Phys. 126, 014101

66. Parrinello, M. and Rahman, A. (1980) Crystal Structure and Pair Potentials: A Molecular-Dynamics Study, Phys. Rev. Lett. 45, 1196–1199

67. Hess, B., Bekker, H., Berendsen, H. J. C., and Fraaije, J. G. E. M. (1997) LINCS: A Linear Constraint Solver for Molecular Simulations, J. Comp. Chem. 18, 1463–1472

68. Darden, T., York, D., and Pedersen, L. (1993) Particle mesh Ewald: An *N*·log(*N*) method for Ewald sums in large systems, J. Chem. Phys. 98, 10089–10092

69. Daura, X., Gademann, K., Jaun, B. Seebach, D. van Gunsteren, W. F., and Mark, A. E. (1999) Peptide Folding: When Simulation Meets Experiment. Angew. Chem. Int. Ed. **38**, 236-240

