## Supplemental Information for "Computational Insights into How Structural Changes in the Hydrophobic Region of ALS-Linked Mutant SOD1 (G93A) Activate Fluctuations of the Electrostatic Loop: Implications for Aggregation"

**Running title:** Structural analysis of hydrophobic region of SOD1 using MD simulation

*To whom correspondence should be addressed.
Prof. Kazuaki Fukushima
Department of Chemistry, School of Medicine, Hyogo Medical University
1-1 Mukogawa-cho, Nishinomiya, Hyogo 663-8501, Japan,


Prof. Noriko Fujiwara
Department of Biochemistry, School of Medicine, Hyogo Medical University
1-1 Mukogawa-cho, Nishinomiya, Hyogo 663-8501, Japan,


S1 Aggregate formation

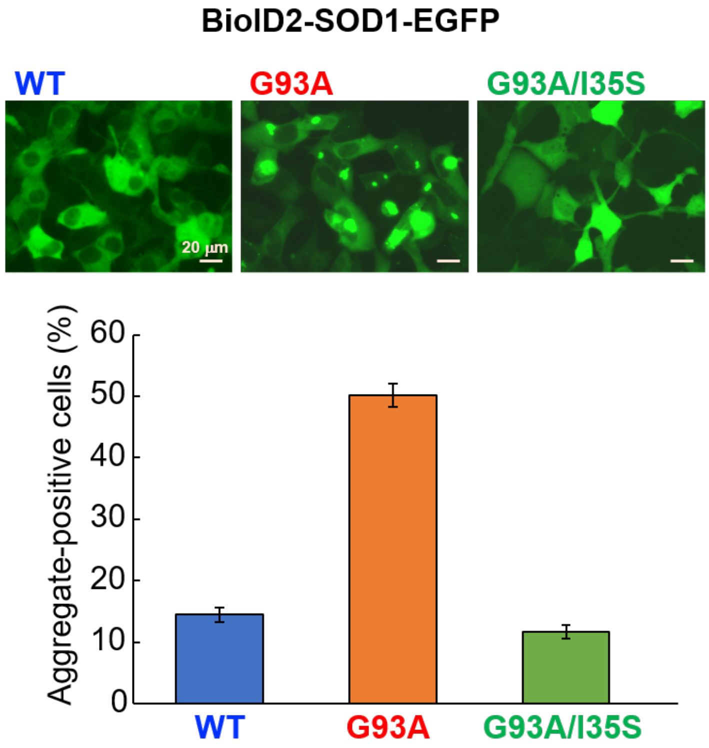


Figure S1. Aggregates formation of BioID2-SOD1-EGFP

Additional substituting Ile35 with serine (I35S) suppressed the aggregate formation of the ALS mutant SOD1 (G93A). (Modified from Asai Y et al. Mol Neurobiol. 2025, 62(2):2023-2038)

S2. RMSD (Root Mean Square Deviation)

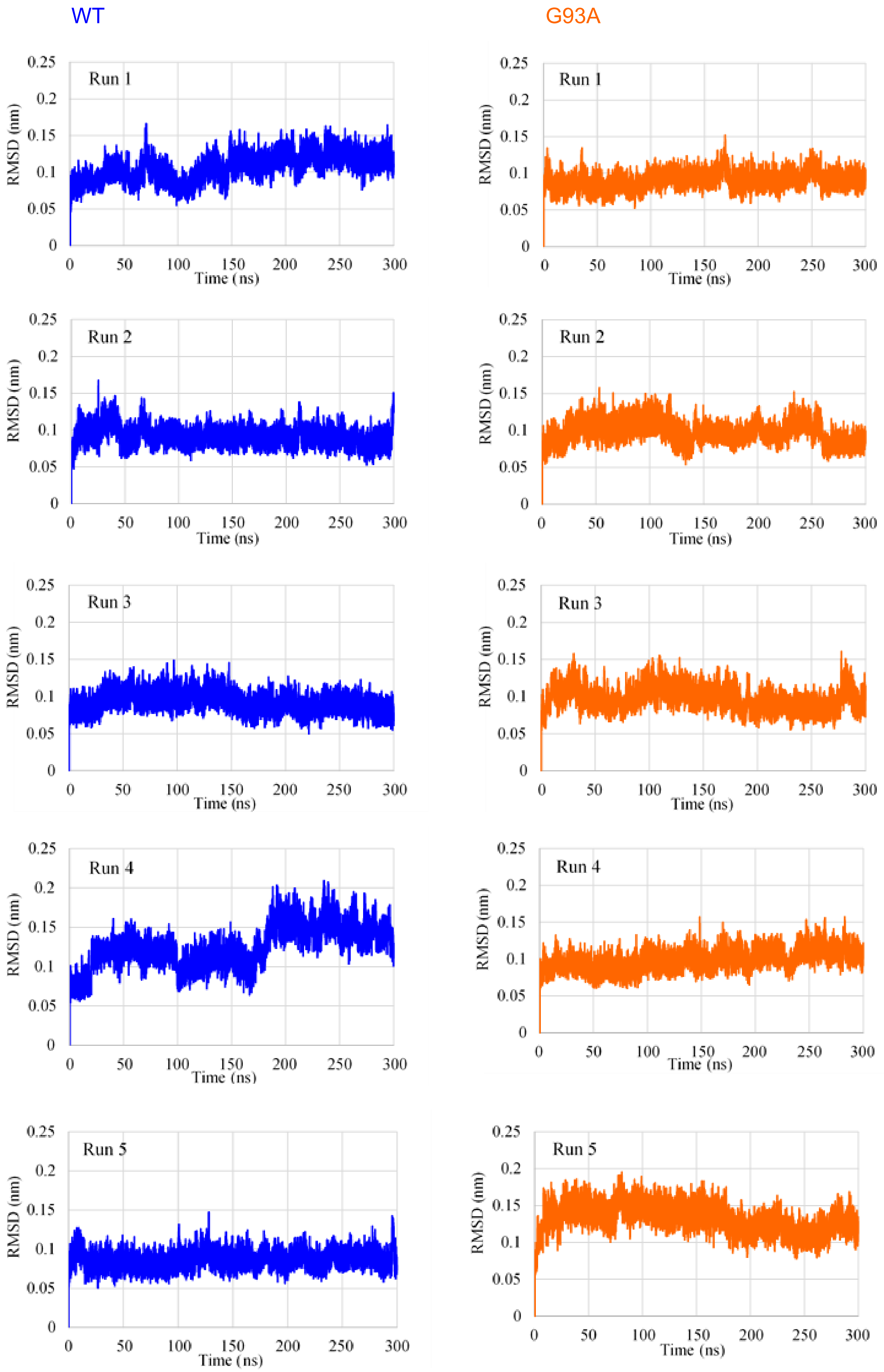


Figure S2. RMSD of WT and G93A.

### S3. RMSF (Root Mean Square Fluctuation)

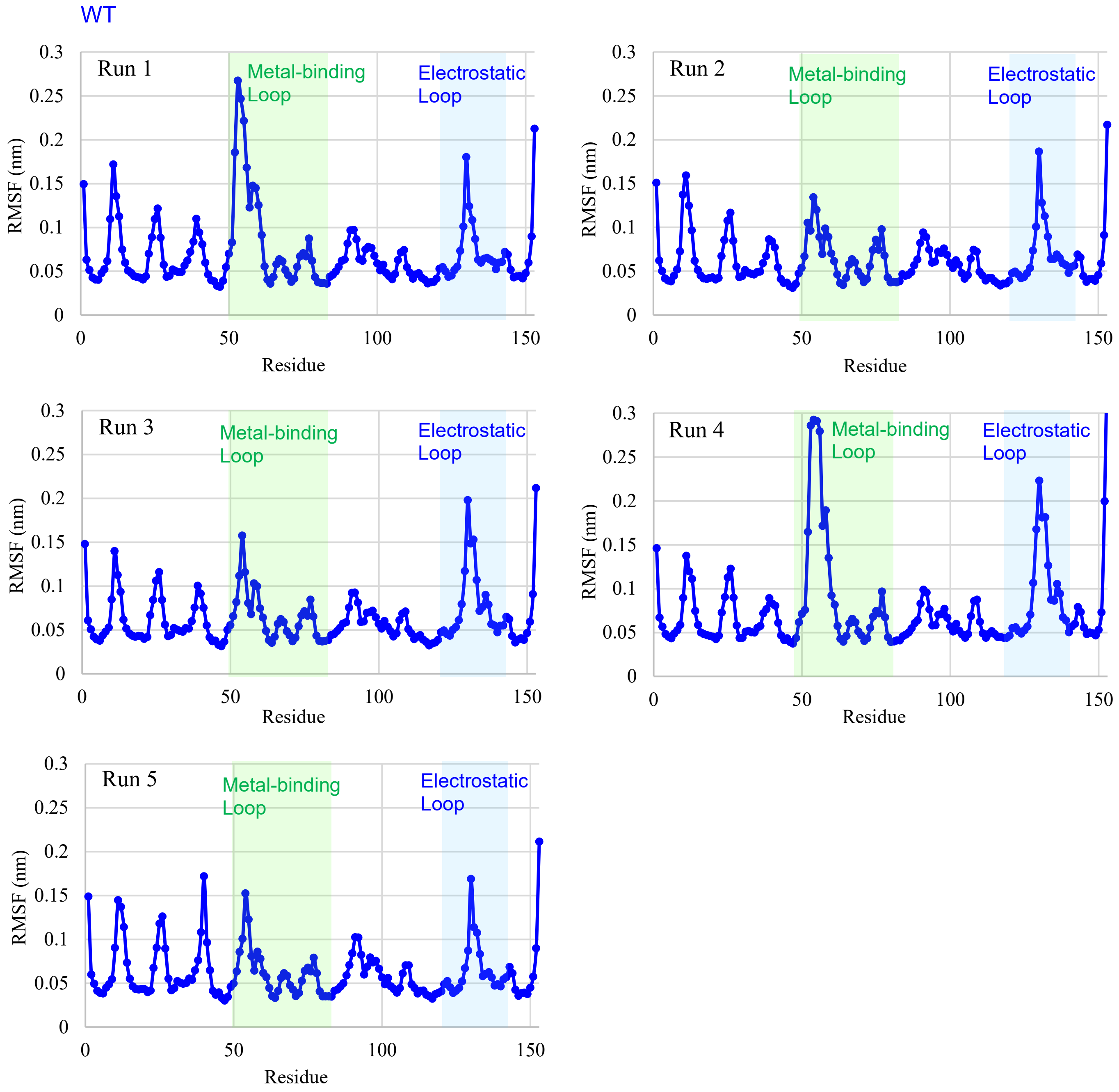


Figure S3-1. RMSF of WT.

The metal-binding loop (49-82) and the electrostatic loop (121-142) are indicated by green and blue bands, respectively.

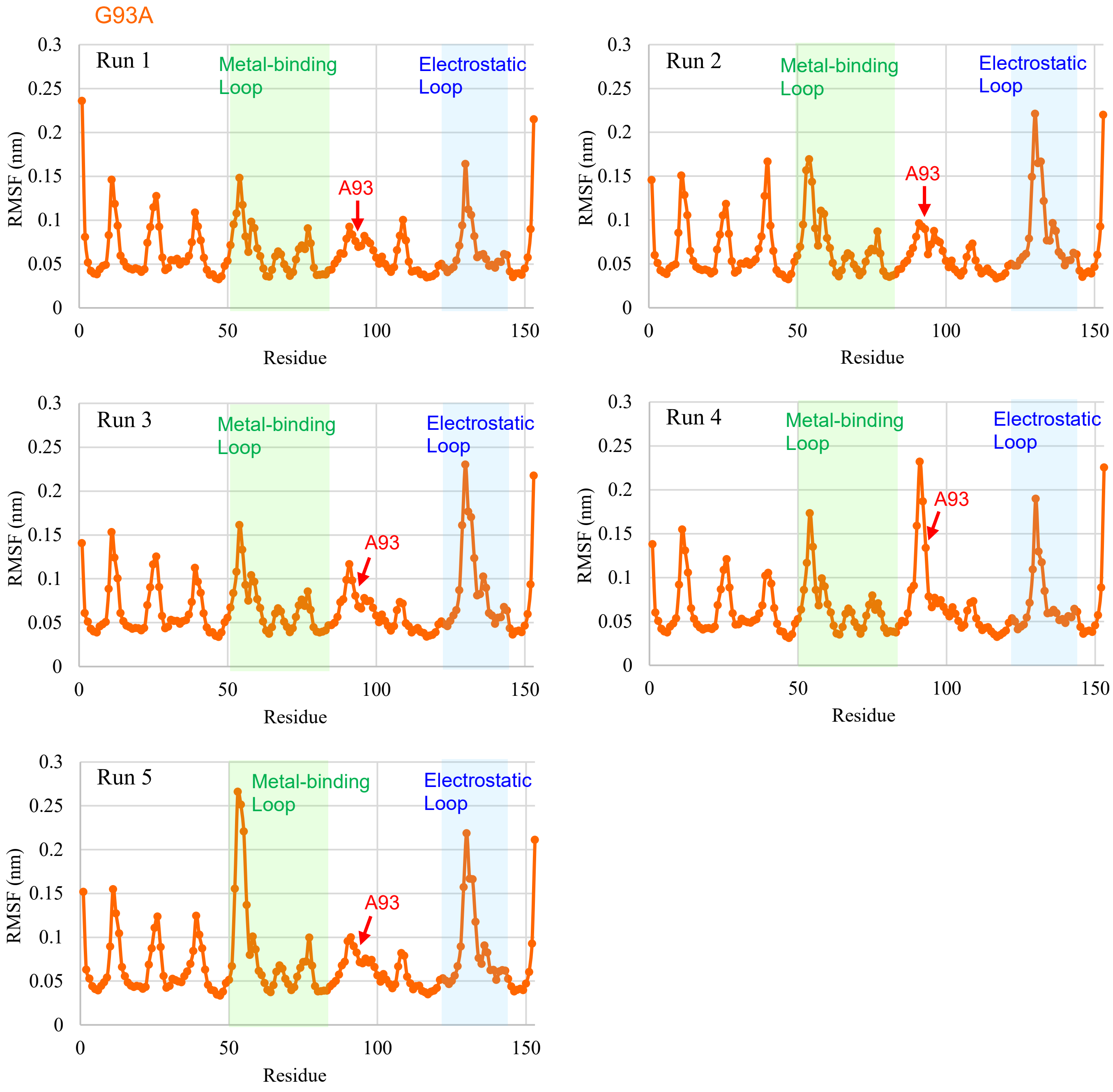


Figure S3-2. RMSF of G93A.
The metal-binding loop (49-82) and the electrostatic loop (121-142) are indicated by green and blue bands, respectively.

Table S3-1. Average RMSF value (nm) of the top of peaks in 5 runs.

| Peak | WT | G93A | difference |
|---|---|---|---|
| *N*-terminus (A1) | 0.14852 | 0.16232 | +0.01380 |
| Loop I (D11) | 0.15048 | 0.15186 | +0.00196 |
| Loop II (N26) | 0.12046 | 0.12314 | +0.00268 |
| Loop III (E40) | 0.10480 | 0.11502 (T39) | +0.01022 |
| Loop IV (T54) | 0.19662 | 0.18062 | -0.01600 |
| Loop V (K91) | 0.09674 | 0.12738 | +0.03064 |
| Loop VI (D109) | 0.07494 | 0.07938 | +0.00444 |
| Loop VII (G130) | 0.19122 | 0.20472 | +0.01350 |
| *C*-terminus (Q153) | 0.24972 | 0.21776 | -0.03196 |

### S4. RMSD of Electrostatic Loop and Metal-binding Loop

#### Electrostatic Loop

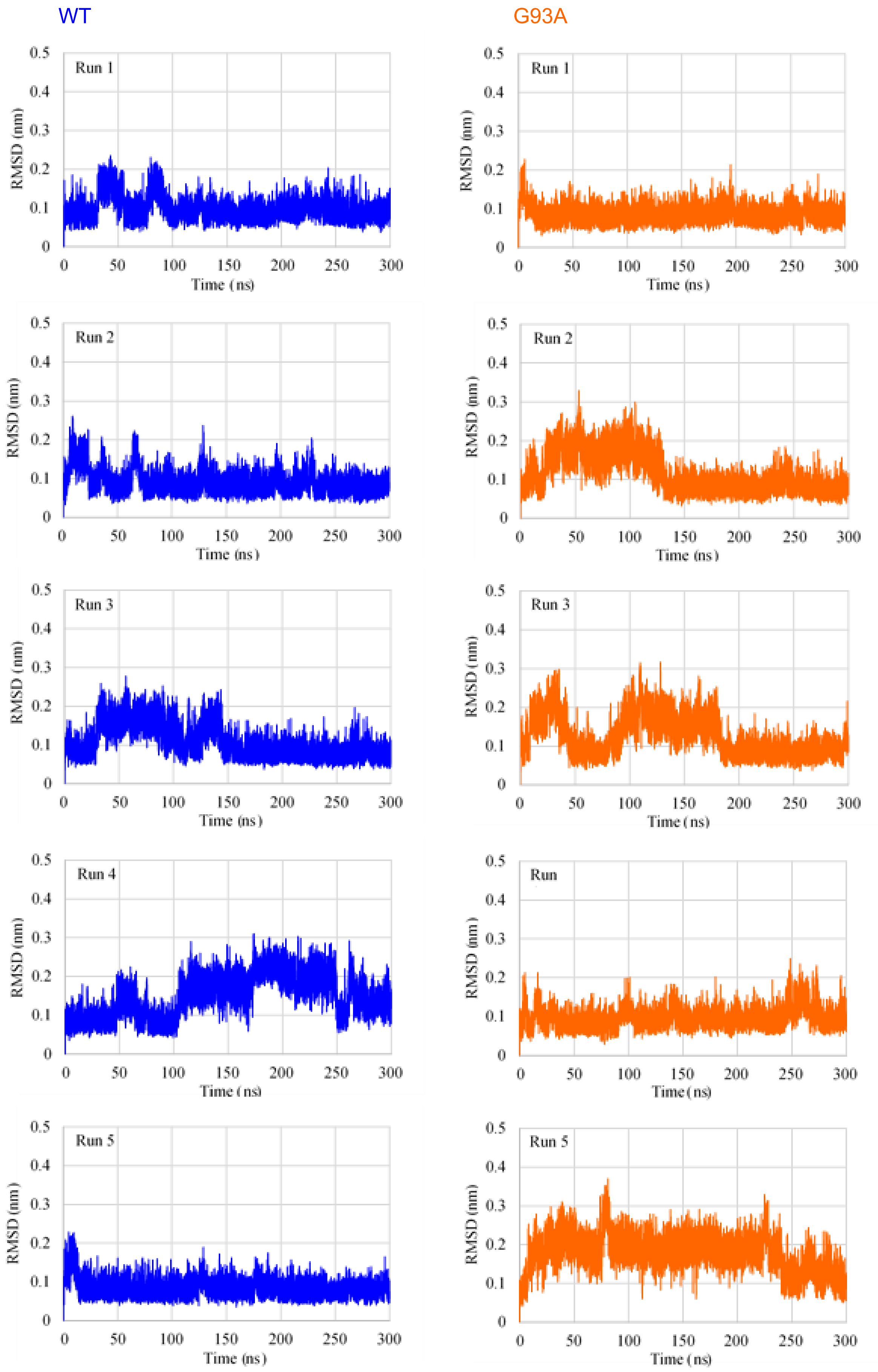


Figure S4-1. RMSD of the electrostatic loop (121-142) of WT and G93A.

Metal-binding Loop

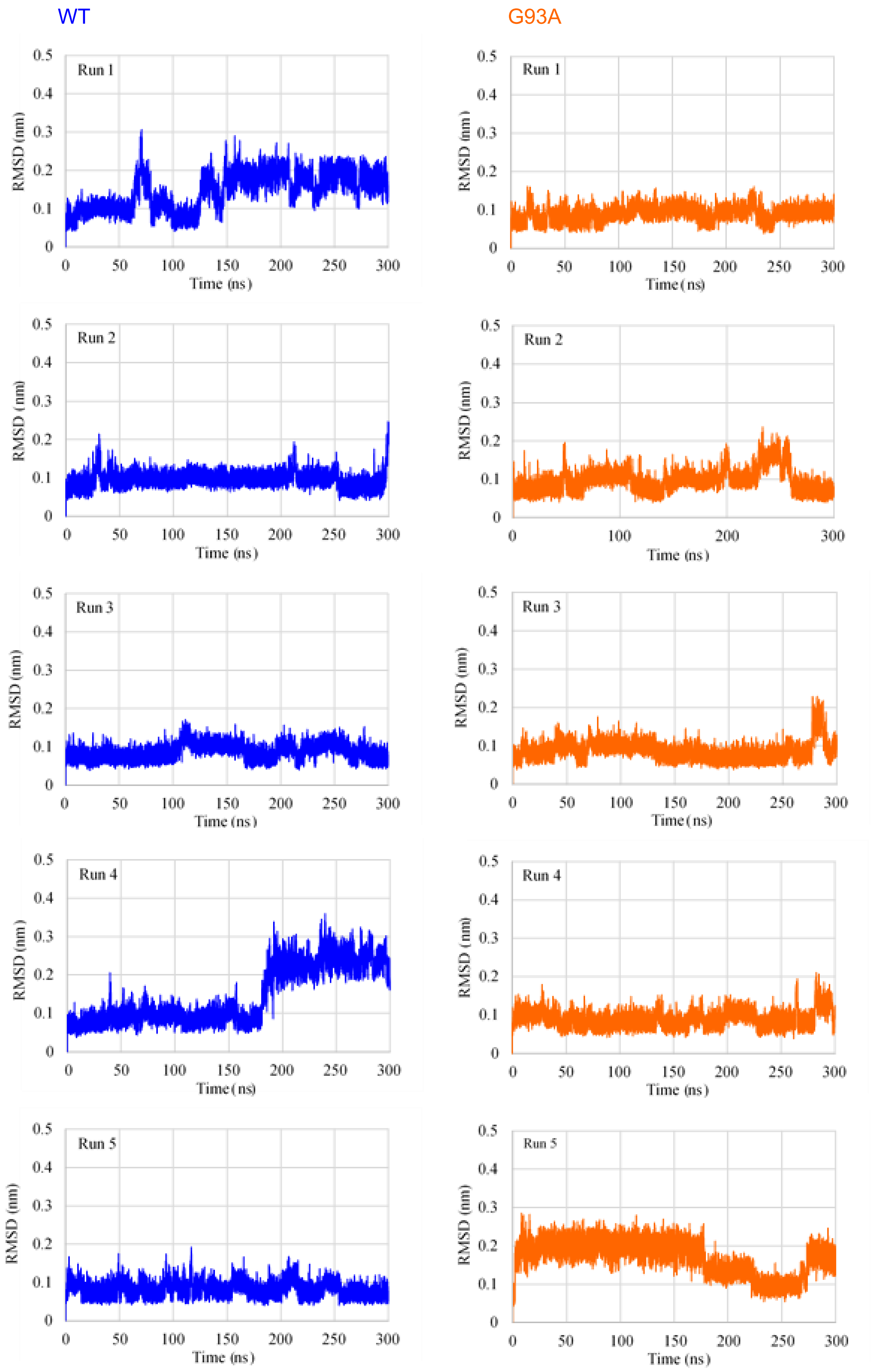


Figure S4-2. RMSD of the metal-binding loop (49-82) of WT and G93A.

Table S4-1. Statistical analysis of RMSD distributions for the metal-binding loop (E49-G82) and electrostatic loop (E121-S142) after least-squares fitting to backbone atoms (C, $C_\alpha$, N).

| Values | Metal-binding Loop | | Electrostatic Loop | |
|---|---|---|---|---|
| | WT | G93A | WT | G93A |
| Sample Number | 150,000 | 150,000 | 150,000 | 150,000 |
| Average | 0.110810 | 0.106180 | 0.105251 | 0.119800 |
| Standard Deviation | 0.0512027 | 0.0429731 | 0.0431646 | 0.0514821 |
| Minimum | 0.0363725 | 0.0376133 | 0.0310182 | 0.0289249 |
| 25 % Percentile | 0.0768494 | 0.0767261 | 0.0749873 | 0.0797214 |
| Median | 0.0944849 | 0.0936537 | 0.0904756 | 0.100708 |
| 75 % Percentile | 0.118951 | 0.117963 | 0.123621 | 0.159163 |
| Maximum | 0.360004 | 0.310182 | 0.285588 | 0.370301 |

### S5. RMSD of β-Strand 8 (L144-I149)

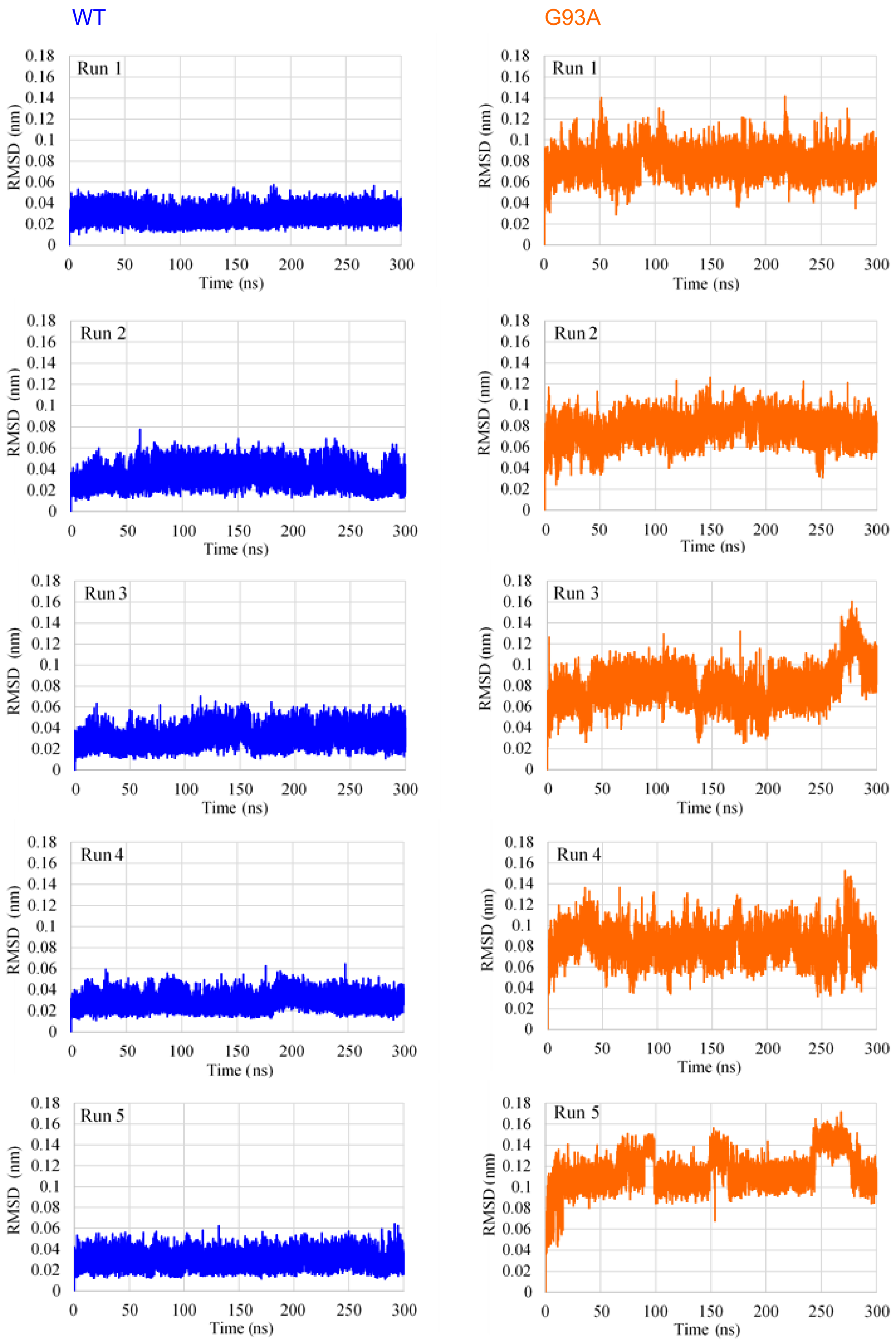


Figure S5. RMSD of β-strand 8 (L144-I149) of WT and G93A.

S6. Distribution of Distances between Cα of G(A)93 and Cα of Proximal Residues

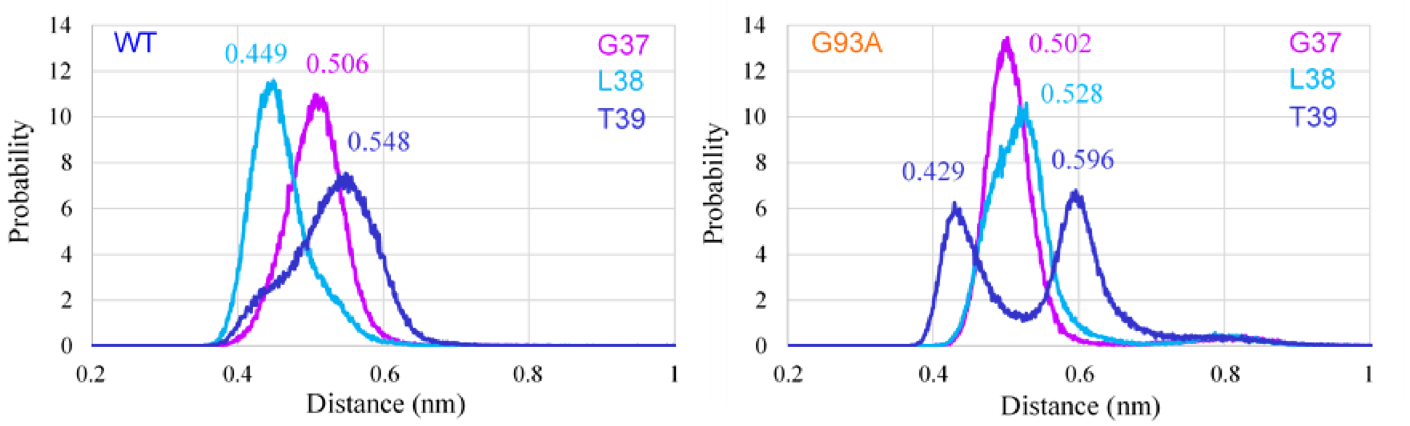


Figure S6-1. Average of five runs for distribution of distances between Cα of G(A)93 and Cα of G37, L38, and T39 in WT and G93A.

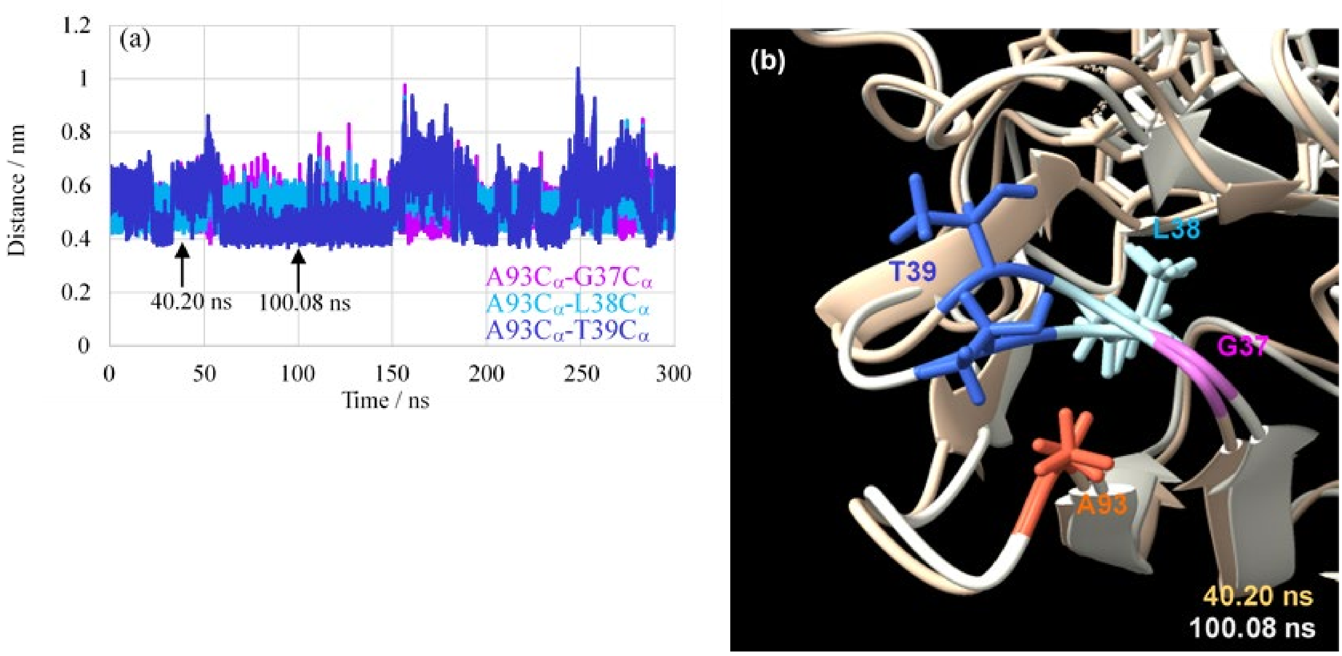


Figure S6-2. (a) Time course of distance between Cα of A93 and Cα of the proximal residues in G93A for Run 1. (b) Typical geometries of T39 at 40.20 and 100.08 ns in the time course of G93A for Run 1.

### S7. Hydrogen Bond between V119 and L144

Distance between V119 CO and L144 NH

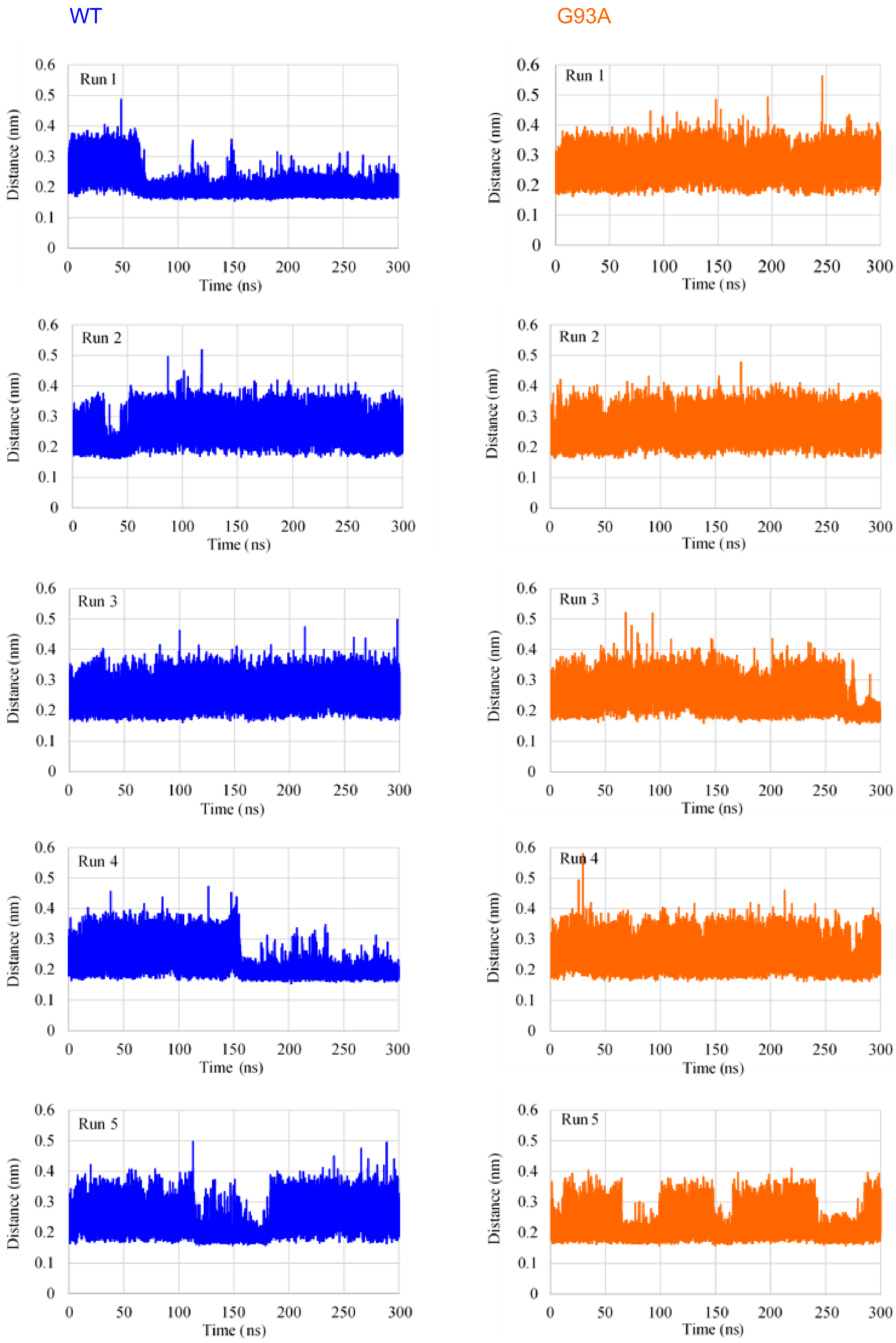


Figure S7. Time course of distance between V119 C=O and L144 N-H of WT and G93A.

### S8. Principal Component Analysis of WT and G93A.

#### WT, Run 1

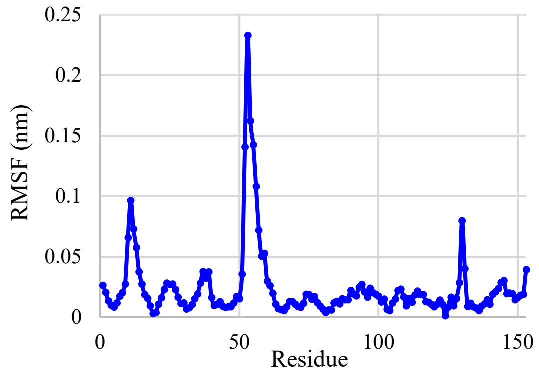


Figure S8-1. RMSF (eigenvectors) of principal component 1 (PC1) in Run 1 of WT simulation.

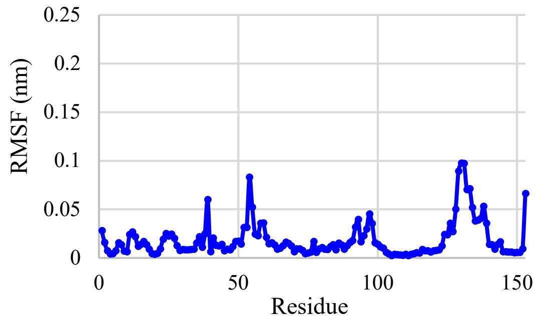


Figure S8-1-1. RMSF (eigenvectors) of principal component 1 (PC1) in Run 1 of WT simulation for 0 – 60 ns.

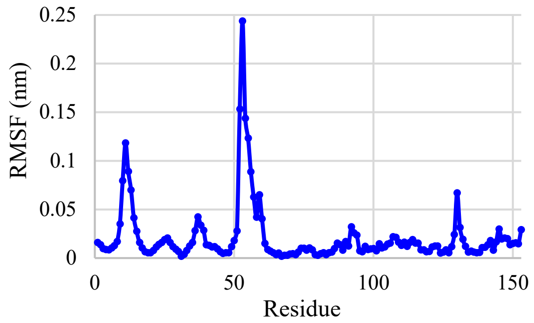


Figure S8-1-2. RMSF (eigenvectors) of principal component 1 (PC1) in Run 1 of WT simulation for 70 – 300 ns.

#### WT, Run 2

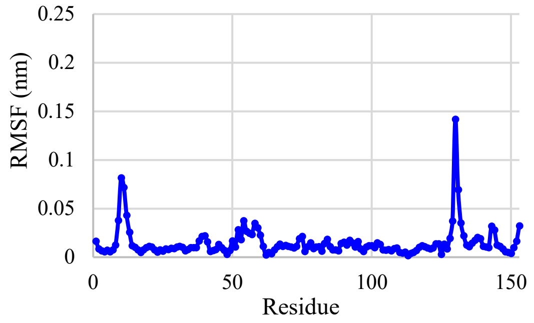


Figure S8-2. RMSF (eigenvectors) of principal component 1 (PC1) in Run 2 of WT simulation.

WT, Run 3

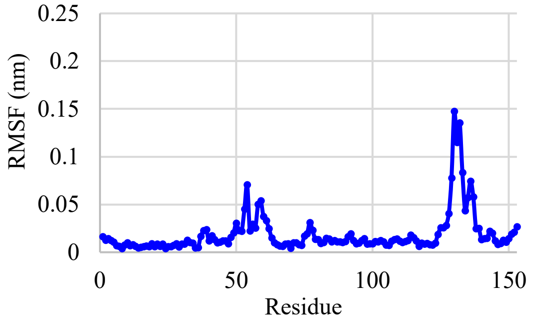


Figure S8-3. RMSF (eigenvectors) of principal component 1 (PC1) in Run 3 of WT simulation.

WT, Run 4

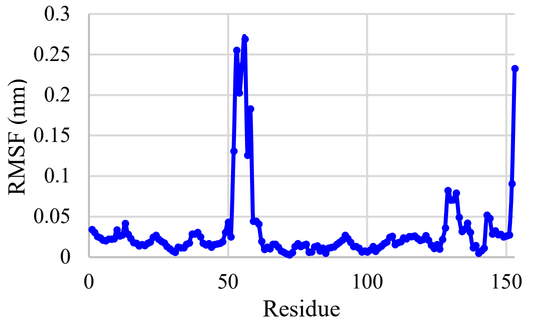


Figure S8-4. RMSF (eigenvectors) of principal component 1 (PC1) in Run 4 of WT simulation.

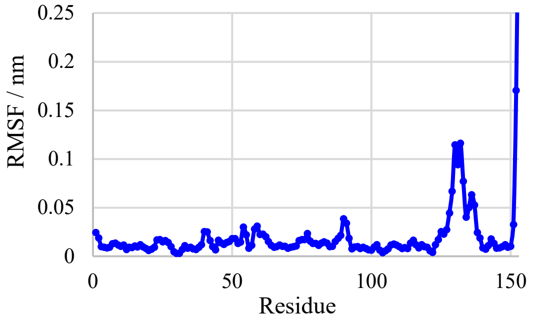


Figure S8-4-1. RMSF (eigenvectors) of principal component 1 (PC1) in Run 4 of WT simulation for 0 – 150 ns.

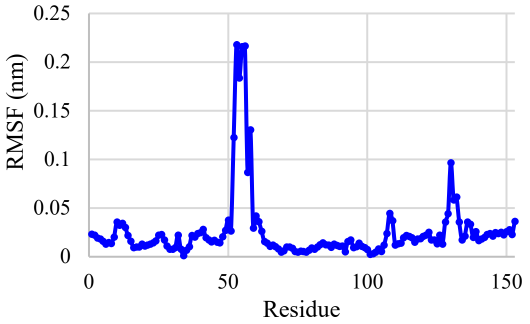


Figure S8-4-2. RMSF (eigenvectors) of principal component 1 (PC1) in Run 4 of WT simulation for 160 – 300 ns.

WT, Run 5

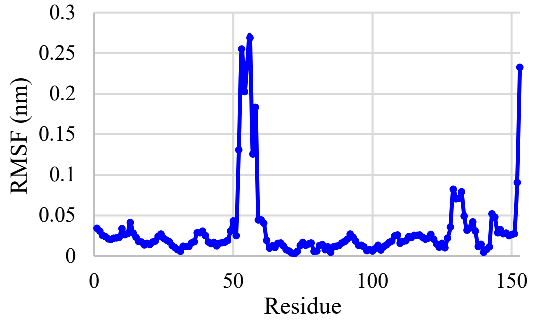


Figure S8-5. RMSF (eigenvectors) of principal component 1 (PC1) in Run 5 of WT simulation.

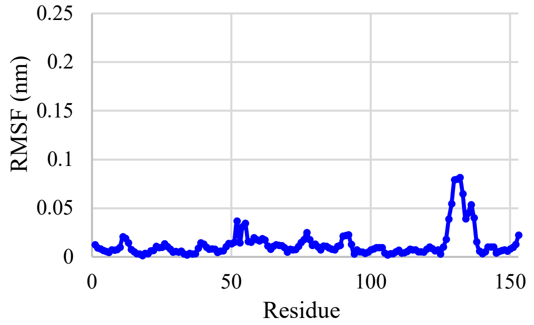


Figure S8-5-1. RMSF (eigenvectors) of principal component 1 (PC1) in Run 5 of WT simulation for 0 – 110 ns.

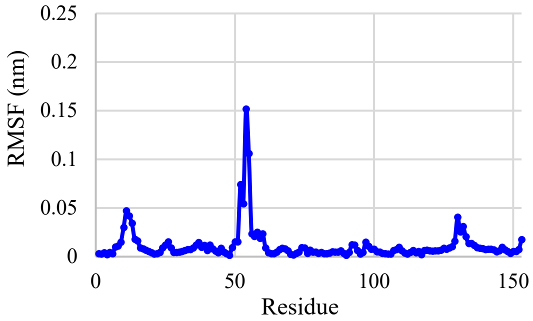


Figure S8-5-2. RMSF (eigenvectors) of principal component 1 (PC1) in Run 5 of WT simulation for 115 – 180 ns.

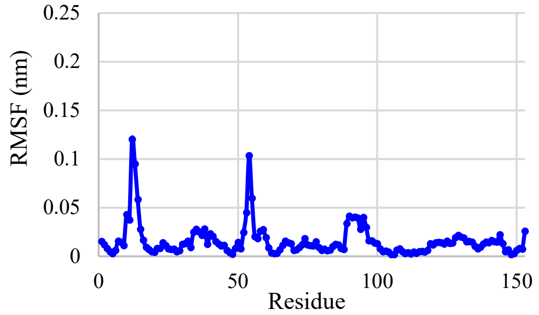


Figure S8-5-3. RMSF (eigenvectors) of principal component 1 (PC1) in Run 5 of WT simulation for 190 – 300 ns.

G93A, Run1

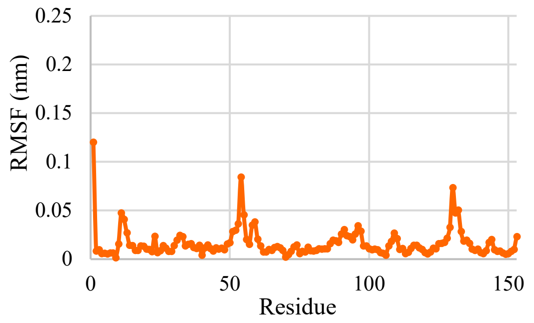


Figure S8-6. RMSF (eigenvectors) of principal component 1 (PC1) in Run 1 of G93A simulation.

G93A, Run 2

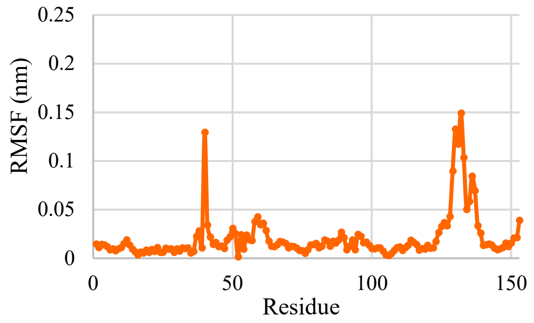


Figure S8-7. RMSF (eigenvectors) of principal component 1 (PC1) in Run 2 of G93A simulation.

G93A, Run 3

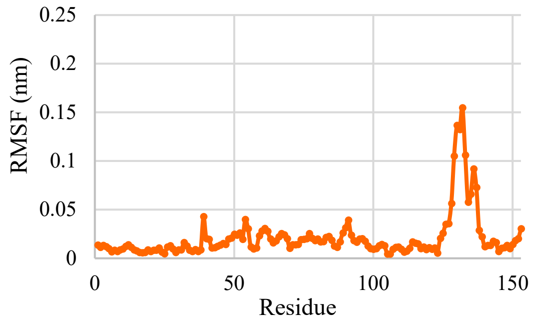


Figure S8-8. RMSF (eigenvectors) of principal component 1 (PC1) in Run 3 of G93A simulation.

G93A, Run 4

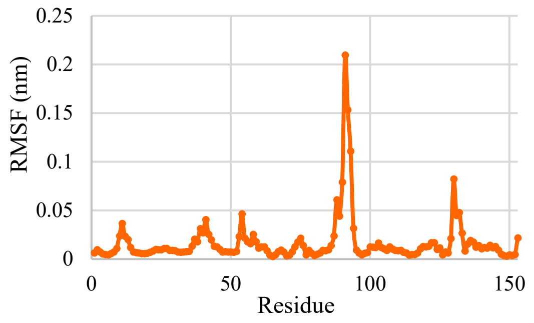


Figure S8-9. RMSF (eigenvectors) of principal component 1 (PC1) in Run 4 of G93A simulation.

G93A, Run 5

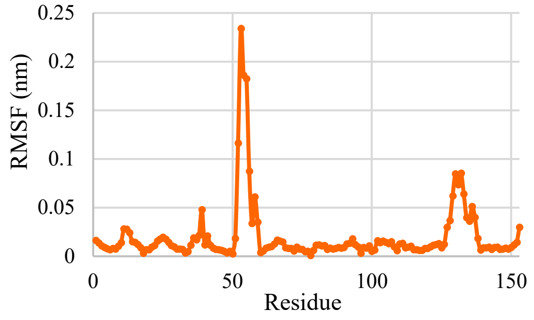


Figure S8-10. RMSF (eigenvectors) of principal component 1 (PC1) in Run 5 of G93A simulation.

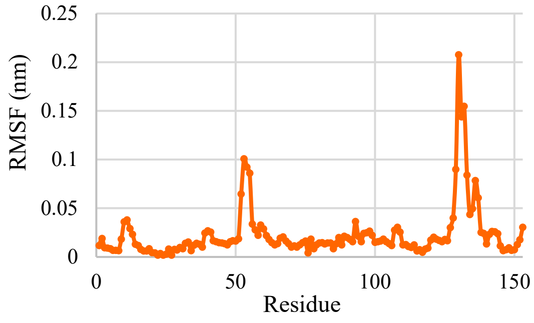


Figure S8-10-1. RMSF (eigenvectors) of principal component 1 (PC1) in Run 5 of G93A simulation for 0–60 ns.

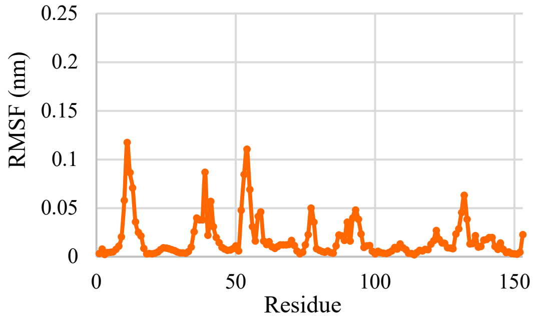


Figure S8-10-2. RMSF (eigenvectors) of principal component 1 (PC1) in Run 5 of G93A simulation for 65–100 ns.

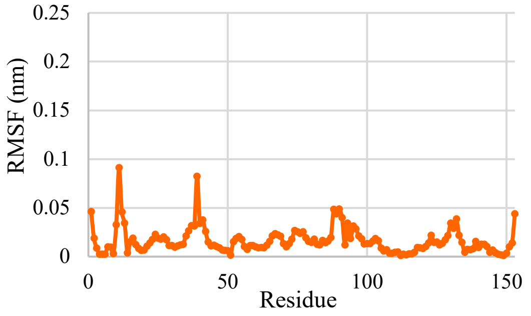


Figure S8-10-3. RMSF (eigenvectors) of principal component 1 (PC1) in Run 5 of G93A simulation for 105–145 ns.

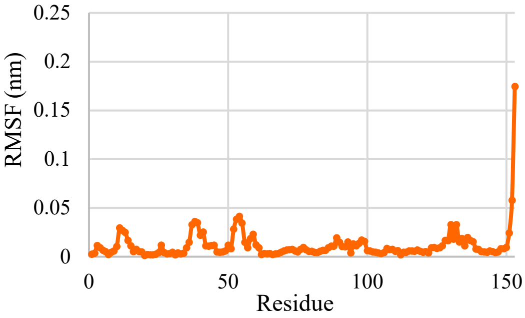


Figure S8-10-4. RMSF (eigenvectors) of principal component 1 (PC1) in Run 5 of G93A simulation for 150–165 ns.

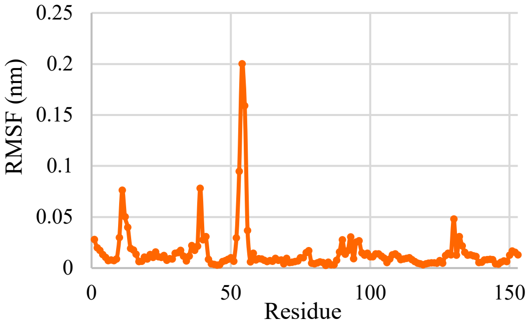


Figure S8-10-5. RMSF (eigenvectors) of principal component 1 (PC1) in Run 5 of G93A simulation for 170–240 ns.

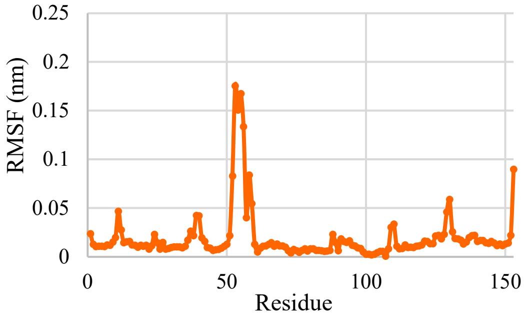


Figure S8-10-6. RMSF (eigenvectors) of principal component 1 (PC1) in Run 5 of G93A simulation for 245–280 ns.

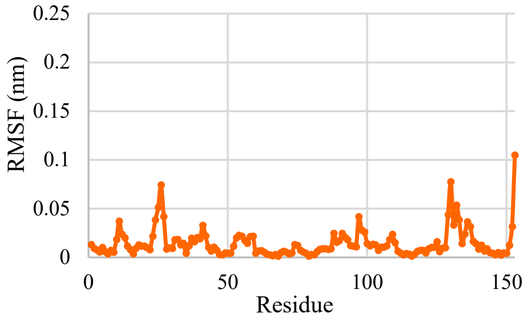


Figure S8-10-7. RMSF (eigenvectors) of principal component 1 (PC1) in Run 5 of G93A simulation for 290–300 ns.

S9. Distance Distribution between L8 and Selected Proximal Residues

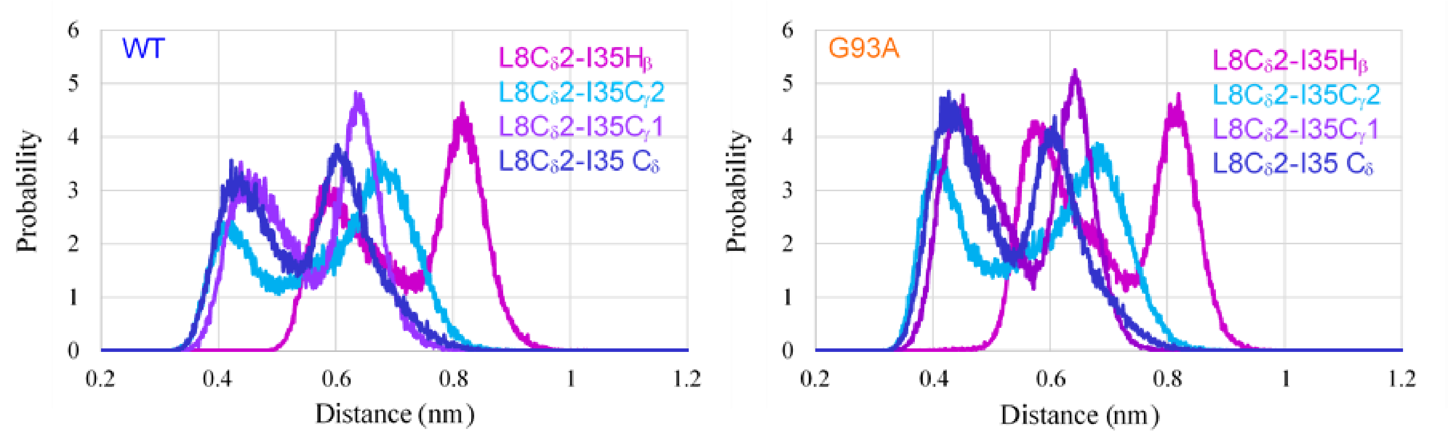


Figure S9-1 Average of five runs for distance distribution between Cδ2 of L8 and atoms of I35 in WT and G93A.

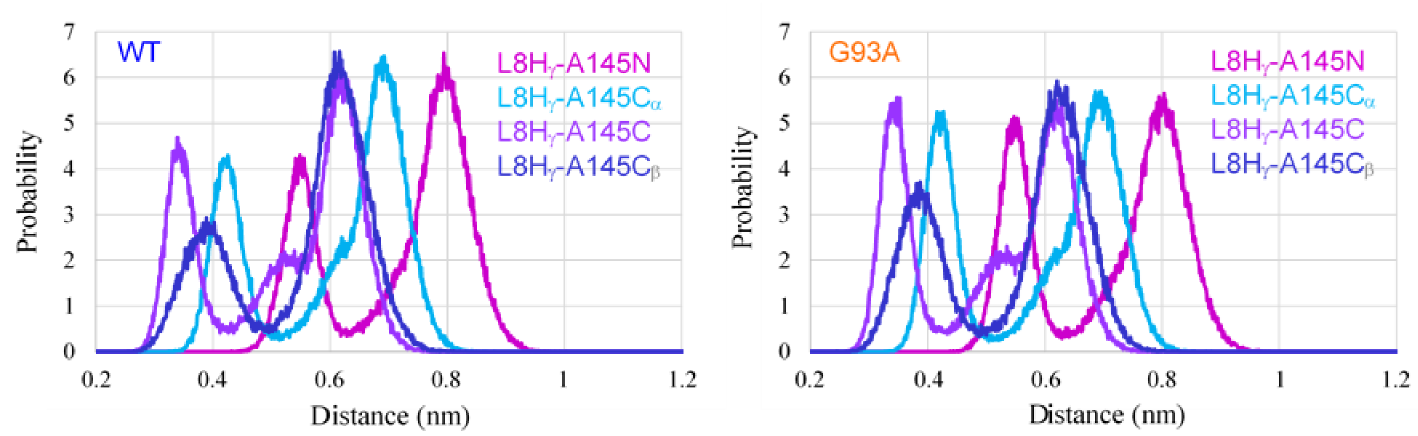


Figure S9-2 Average of five runs for distance distribution between Hγ of L8 and atoms of A145 in WT and G93A.

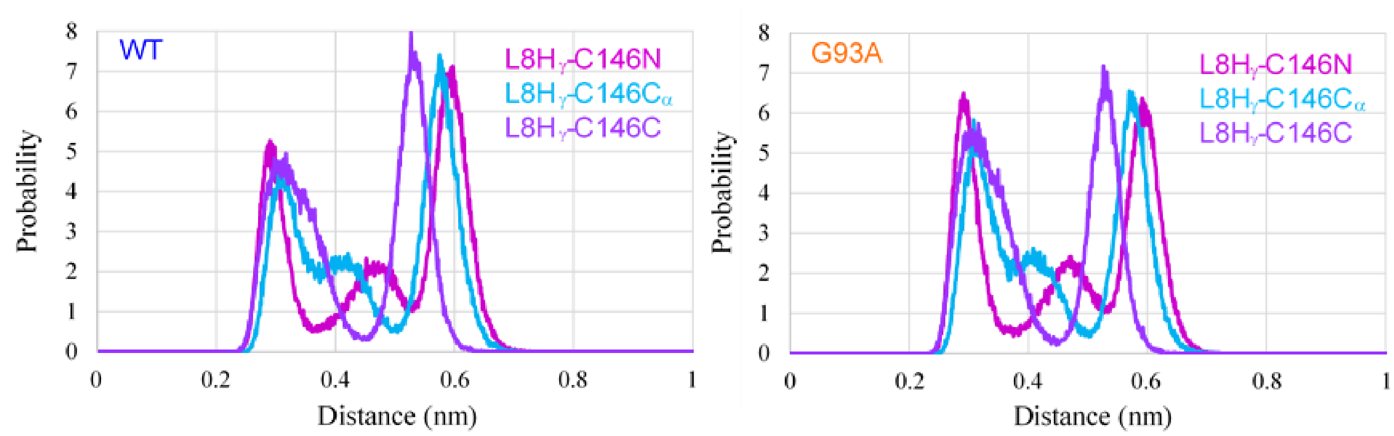


Figure S9-3 Average of five runs for distance distribution between Hγ of L8 and atoms of C146 in WT and G93A.

The distance distribution in Figs. S9-1 to S9-3 showed two large peaks. The short-distance peaks between $L8C_\delta$ and the atoms of I35 mean that $L8C_\delta$ points toward I35, and the short-distance peaks between $L8H_\gamma$ and the atoms of A145 and C146 mean that $L8H_\gamma$ points toward the β-strand 8. The height of the short-distance peaks is lower than that of the long-distance peaks in WT, whereas in G93A, the heights of both peaks are comparable. This means that the ratio of the conformation in which $L8H_\gamma$ points toward the β-strand 8 is larger in G93A than in WT.

S10. Influence of G93A/I35S double mutation

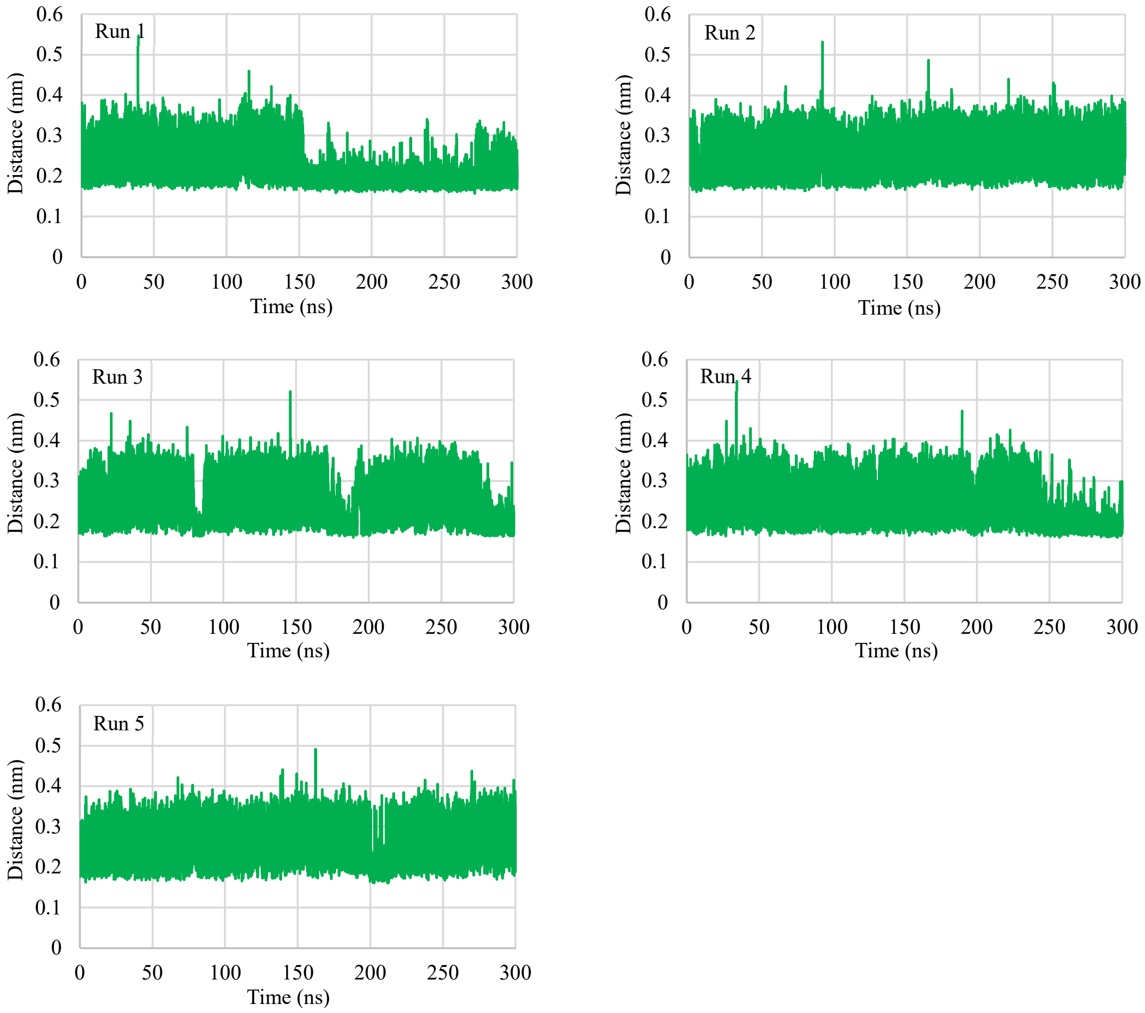


Figure S10-1. Time Course of Hydrogen-bond Formation between V119 C=O and L144 N-H in G93A/I35S.

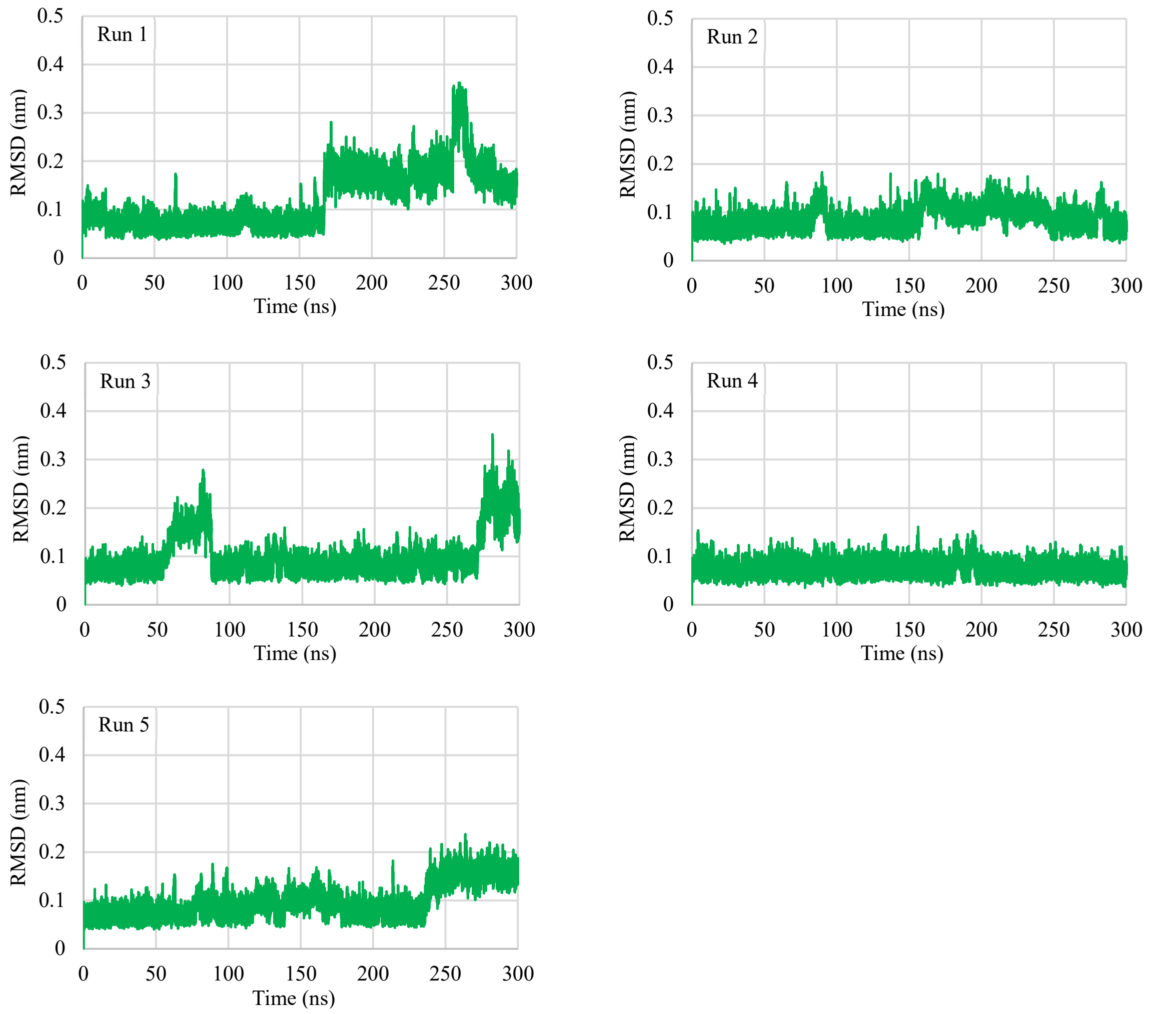


Figure S10-2. RMSD of the Metal-binding Loop (49-82) in G93A/I35S.

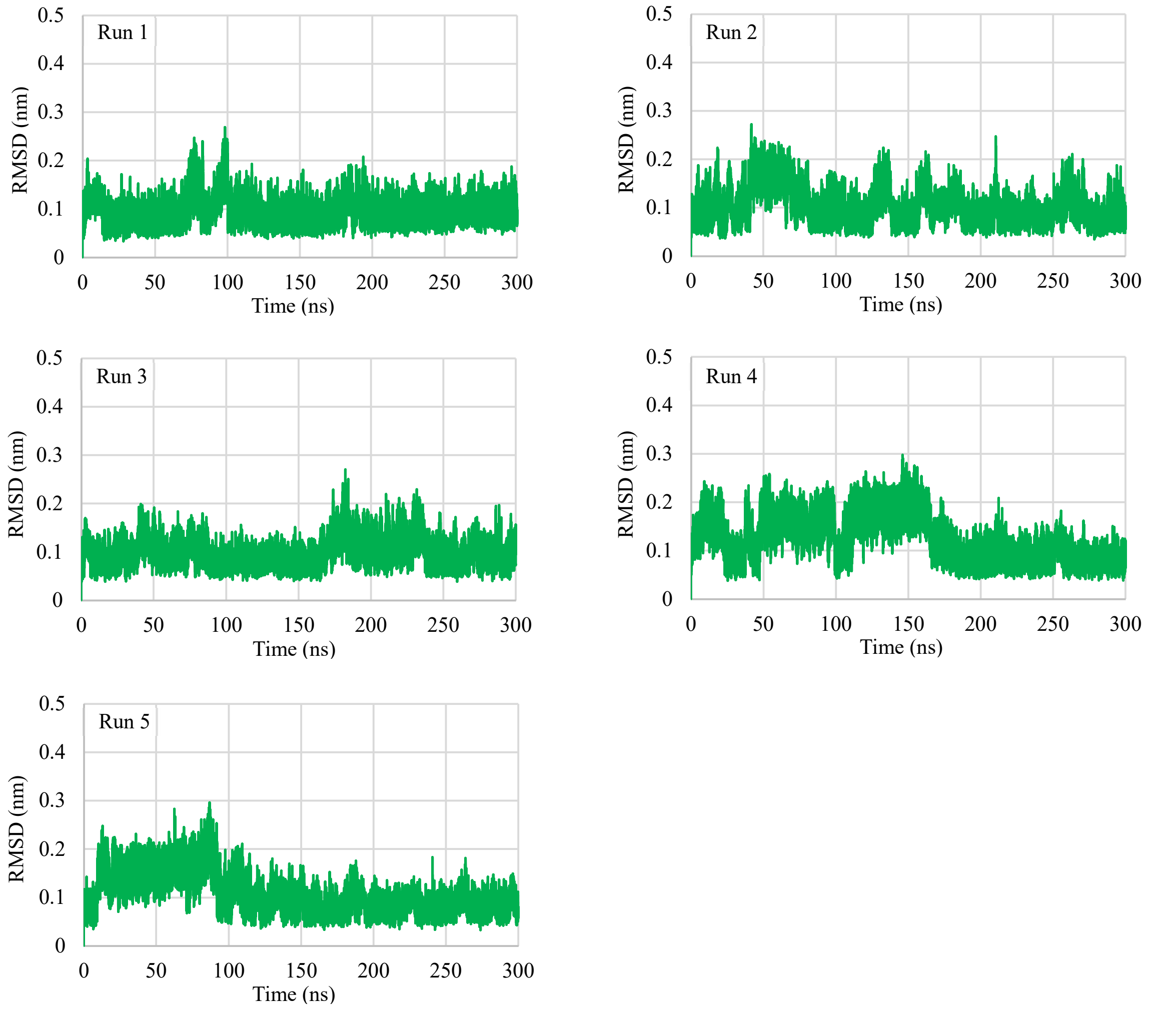


Figure S10-3. RMSD of Electrostatic Loop (121-142) in G93A/I35S.

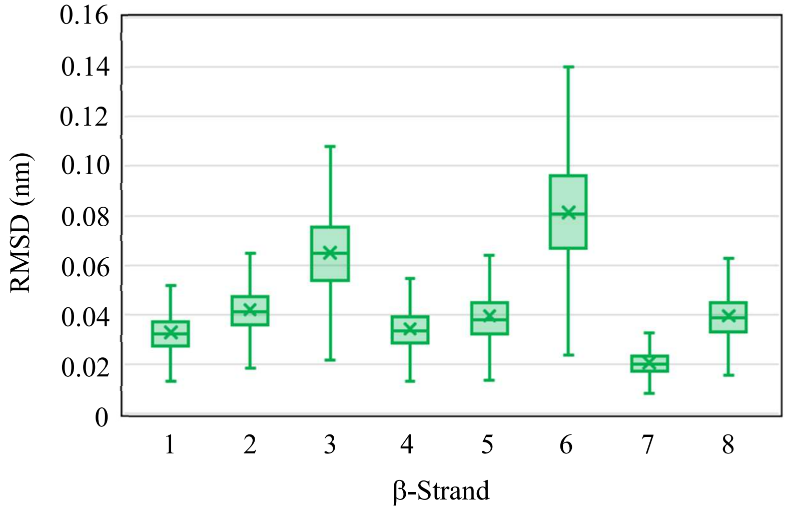


Figure S10-4. Box-and-whisker plot of RMSD of the β-strand 1 to 8 of G93A/I35S. The box represents IQR (interquartile range: 25-75th percentile), the horizontal line inside the box represents the median value, and the cross mark represents the average value. The top of the whiskers above the box represents the maximum value, and that below the bottom represents the minimum value.

S11. Hydrogen-bonding network involved in the fluctuation of the electrostatic loop

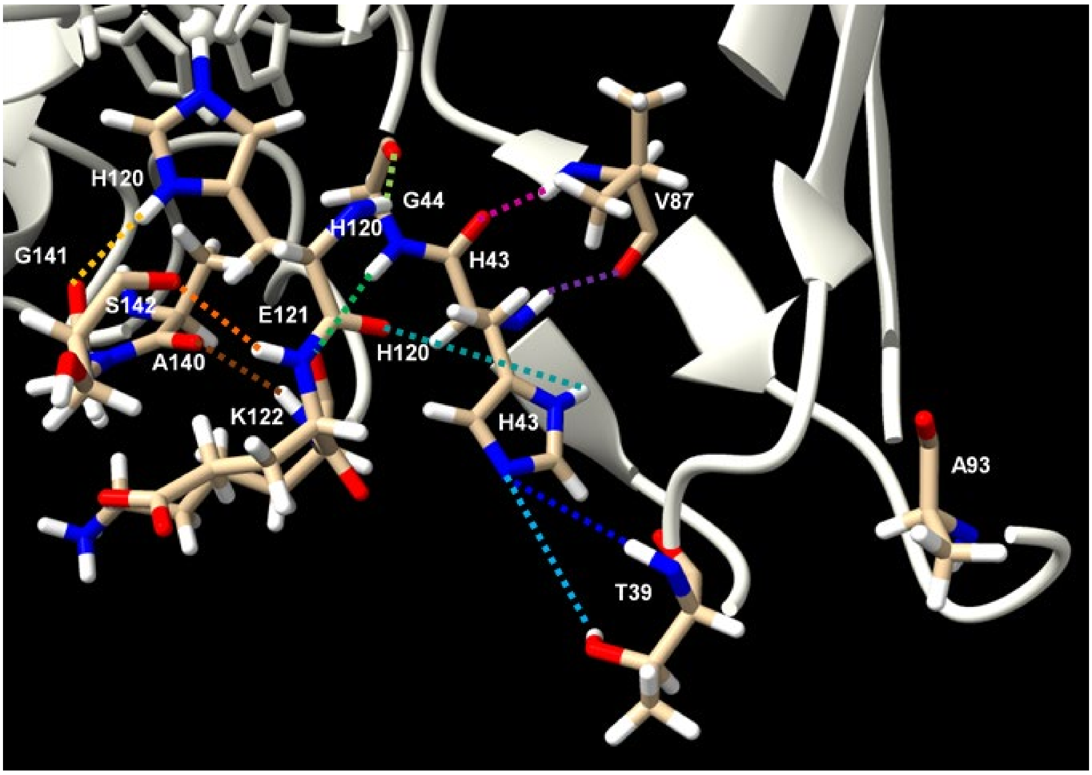


Figure S11-1. Hydrogen bond network of T39, H43, V87, G44, H120, E121, K122, A140, G141, and S142.

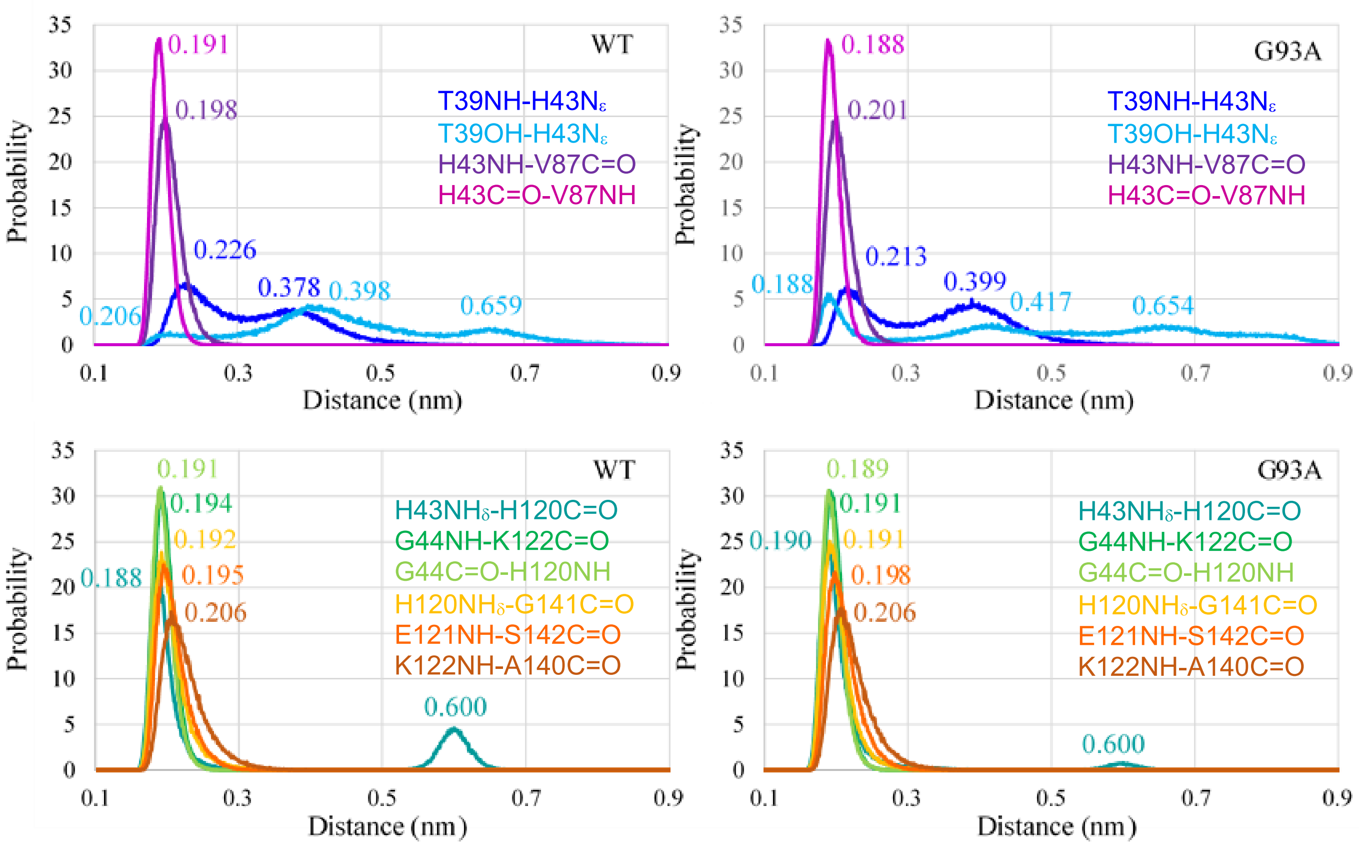


Figure S11-2. Average of five runs for distance distribution of hydrogen bond in electrostatic loop and related residues in WT and G93A.

S12. Comparison of X-ray Structures

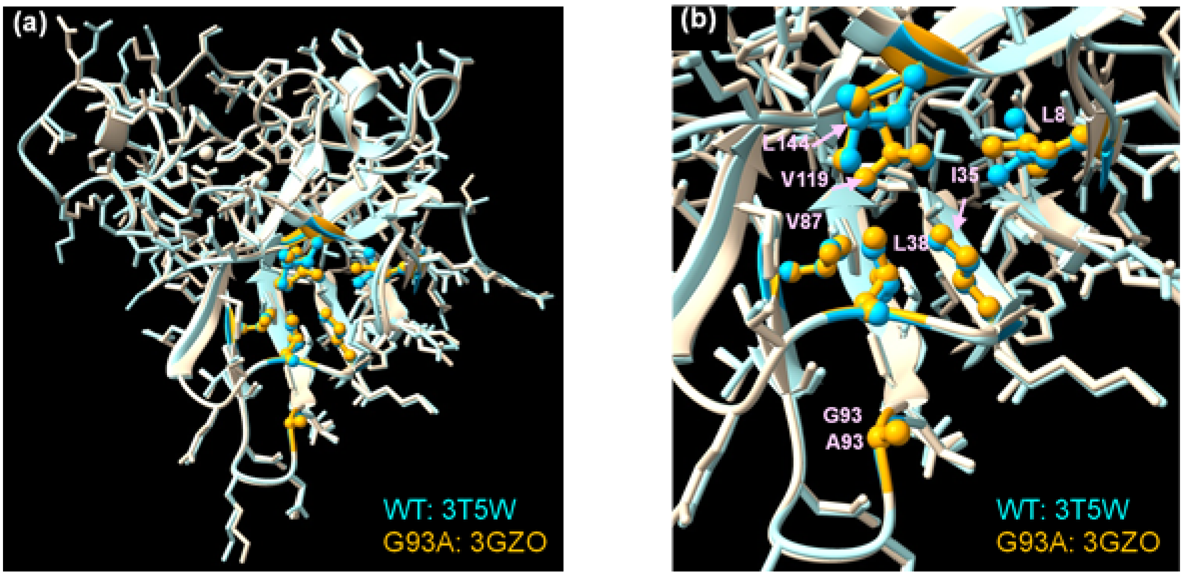


Figure S12. X-ray structures of WT (3T5W) and G93A (3GZO).
(a) Whole structures of monomer proteins, (b) Hydrophobic region.

Table S12. Interatomic distances (nm)

| | WT | G93A | Difference (G93A-WT) |
|---|---|---|---|
| G(A)93C=O – L38 N | 0.2970 | 0.3045 | +0.0075 |
| G(A)93C=O – G37 N | 0.3223 | 0.2969 | -0.0254 |
| V119 C=O – L144 N | 0.2939 | 0.2942 | +0.0003 |
| L38 $C_\alpha$ – I35 $C_\alpha$ | 0.7453 | 0.7330 | -0.0120 |
| L38 $C_\alpha$ – L144 $C_\alpha$ | 0.8660 | 0.8605 | -0.0055 |
| L38 $C_\alpha$ – V87 $C_\alpha$ | 1.0236 | 1.0061 | -0.0175 |
